# Enhanced Pathogenicity and Contact Transmissibility of Human-origin Avian Influenza H5N1 Clade 2.3.4.4b Genotype B3.13 Compared to D1.1 in Ferrets

**DOI:** 10.64898/2026.08.10.744032

**Authors:** Ahmed M. Elsayed, Ramya S. Barre, Mahmoud Bayoumi, Alvaro Padron, Hossein Batebi, Vinay Shivanna, Roy N. Platt, Fiona Burmeister, Joshua Castro, Arash Rahmani, Juliane Lang, Chengjin Ye, Timothy J. C. Anderson, Roland Netz, Aitor Nogales, Robert P. de Vries, Geert-Jan Boons, Adolfo García-Sastre, Elsayed M. Abdelwhab, Gregory C. Ippolito, Luis Martinez-Sobrido

**Affiliations:** Host-pathogen interactions (HPI) and Disease Intervention and Prevention (DIP) programs, Texas Biomedical Research Institute, San Antonio, TX 78227, USA; Center of Scientific Excellence for Influenza Viruses, National Research Centre, Giza, Egypt; Virology Department, Faculty of Veterinary Medicine, Cairo University, Giza, Egypt; Department of Physics, Freie Universität Berlin, Berlin, Germany; Institute of Molecular Virology and Cell Biology, Friedrich-Loeffler-Institut, Federal Research Institute for Animal Health, Greifswald-Insel Riems, Germany; Center for Animal Health Research, CISA-INIA-CSIC, Madrid, Spain; Center for Influenza Disease and Emergence Response (CIDER), Madrid, Spain; Division of Chemical Biology & Drug Discovery, Utrecht Institute for Pharmaceutical Sciences, Utrecht University, Utrecht, The Netherlands; Complex Carbohydrate and Research Center, University of Georgia, Athens, GA 30602, USA; Department of Chemistry, University of Georgia, Athens, GA 30602, USA; Center for Influenza Disease and Emergence Response (CIDER), Athens GA, USA; Department of Microbiology, Icahn School of Medicine at Mount Sinai, New York, NY 10029, USA; Global Health and Emerging Pathogens Institute, Icahn School of Medicine at Mount Sinai, New York, NY 10029, USA; Department of Medicine, Division of Infectious Diseases, Icahn School of Medicine at Mount Sinai, New York, NY 10029, USA; The Tisch Cancer Institute, Icahn School of Medicine at Mount Sinai, New York, NY 10029, USA; Department of Pathology, Molecular and Cell-Based Medicine, Icahn School of Medicine at Mount Sinai, New York, NY 10029, USA; The Icahn Genomics Institute, Icahn School of Medicine at Mount Sinai, New York, NY 10029, USA

**Keywords:** H5N1 2.3.4.4b, B3.13, D1.1, avian influenza, ferret, infection, pathogenicity, transmission

## Abstract

Since its emergence in 2020, multiple genotypes of the H5N1 clade 2.3.4.4b have been identified, with B3.13 and D1.1 emerging in the USA as two major and concerning genotypes. However, their relative pathogenicity and transmissibility in mammals have not been fully elucidated. We compared the pathogenicity and transmissibility of the first two human H5N1 clade 2.3.4.4b cases caused by B3.13 in Texas (A/Texas/37/2024; HPhTX B3.13) and D1.1 in Louisiana (A/Louisiana/12/2024; HPhLA D1.1) in a ferret model of infection and transmission. HPhTX B3.13 infection resulted in more severe clinical disease and enhanced viral shedding, with evidence of increased transmission relative to HPhLA D1.1. Histopathological analysis revealed more extensive lung pathology in animals infected with HPhTX B3.13, consistent with increased viral loads and inflammatory responses. Importantly, both genotypes showed no significant differences in reactivity to ferret sera raised against candidate vaccine virus (CVV) strains, receptor binding properties, or neuraminidase (NA) activity and thermostability. Whole-genome sequencing revealed no adaptive mutations in HPhTX B3.13 following infection or transmission. In contrast, HPhLA D1.1 showed rapid acquisition of the mammalian-adaptive mutation E627K in infected ferrets and both E627K and Q194K in the only fatal contact animal. Both mutations were associated with enhanced polymerase activity and computational analyses suggested that they enhance interactions with the mammalian host factors ANP32A and B. Our findings indicate that B3.13 is already well adapted for mammalian infection and transmission whereas D1.1 retains evolutionary potential through the rapid acquisition of adaptive mutations, highlighting important genotype-specific differences relevant to zoonotic risk assessment and pandemic preparedness.

**Significance:** Influenza H5N1 viruses continue to diversify genetically while expanding into mammalian hosts, increasing opportunities for viral adaptation and zoonotic transmission, including humans. However, whether the predominant clade 2.3.4.4b genotype differs in its capacity to infect, transmit, and evolve in mammals remains poorly understood. Using the ferret model of influenza infection and transmission, we demonstrated that the currently circulating B3.13 and D1.1 genotypes exhibit distinct pathogenic and transmission characteristics despite retaining similar receptor-binding characteristics, NA functions, and antigenic profiles. While B3.13 readily infects and transmits in ferrets and does not develop further adaptive mutations associated with increased replication and transmission, D1.1 rapidly acquires mammalian-adaptive mutations after a single infection and/or transmission event, highlighting its evolutionary potential. These findings show that genotype-specific biological properties can influence zoonotic risk independently of antigenic similarity and emphasize the importance of integrating phenotypic characterization with genomic surveillance to improve pandemic preparedness and guide public health risk assessment.

## Introduction

Influenza A viruses (IAVs) pose a significant threat to global public health, particularly considering the recent outbreaks of avian influenza viruses (AIVs) with zoonotic potential^1^. AIVs primarily circulate within wild birds and poultry but have increasingly demonstrated the ability to infect a wide range of mammalian hosts, including humans ^1,2^. AIVs are classified as high pathogenic (HPAIV) or low pathogenic (LPAIV) based on their molecular characteristics and their ability to cause disease in chickens. HPAIVs are typically associated with the H5 and H7 hemagglutinin (HA) subtypes ^3^. Alarmingly, outbreaks of HPAIV H5N1 are becoming more frequent, and recent surveillance has identified infections in multiple mammalian species including minks, cats, seals, and cattle ^2^. These events highlight the ongoing adaptation of AIVs to mammalian hosts and underscore their spillover potential. Understanding the molecular determinants that govern viral pathogenicity and transmission in mammalian hosts is critical for assessing the spillover potential of these emerging AIVs.

HPAIV H5N1 emerged in 1997 and a total of 993 laboratory-confirmed human cases, including 477 deaths, have been reported from 2003 to 2026, with a lethality rate >48% ^4^. Notably, infections caused by HPAIV H5N1 are associated with high morbidity and mortality rates and have been reported in several mammalian species ^2,5,6^. In 2024, a new genotype (B3.13) of the H5N1 clade 2.3.4.4b emerged in the United States of America (USA) to infect birds and dairy cattle, marking the first known outbreak of H5N1 in ruminants ^2^. Shortly after, a human case was reported in a dairy farm worker following exposure to infected cattle, representing the first documented case of cattle-to-human transmission of HPAIV H5N1 ^7^. Unexpectedly, the H5N1 clade 2.3.4.4b genotype B3.13 resulted in several outbreaks on cattle farms in the USA, with multiple cases of cattle-to-human transmission ^8^. Amid a yearlong outbreak of the H5N1 clade 2.3.4.4b genotype B3.13 on cattle farms, a second genotype, D1.1, was identified in late 2024 in birds and contact human cases ^2^. Subsequently, in early 2025, the D1.1 genotype was further transmitted from birds to cattle, raising additional concerns regarding its cross-species transmission and adaptation potential ^1,9^.

Infection outcomes of the B3.13 H5N1 genotype appear to differ markedly among mammalian hosts ^7,10–13^. Following the first reported human case in Texas in March 2024, B3.13 infections in humans have been primarily associated with conjunctivitis and/or mild respiratory illness, whereas repeated fatal infections have been documented in cats following ingestion of infected milk ^6^. Experimentally, infection in mice was also shown to be associated with high pathogenicity, as reflected by a mean mouse lethal dose (MLD₅₀) of less than 10 plaque forming units (PFU) ^11,12^. These data indicate that host susceptibility and disease severity associated with the B3.13 genotype vary substantially among mammalian species, underscoring the complex host-specific pathogenic potential of the H5N1 virus. In contrast, most human infections associated with the D1.1 genotype have resulted in severe-to-fatal disease outcomes ^14,15^, at least in one case associated with the presence of autoantibodies neutralizing type I interferons (IFNs) ^16^, whereas experimental animal models did not consistently exhibit comparable levels of pathogenicity ^17,18^.

The ferret model remains a cornerstone for evaluating mammalian infection and transmissibility of H5N1 and other influenza viruses and for guiding pandemic risk assessment ^19^. Ferrets are considered one of the best animal models for human influenza research owing to the close similarity of their respiratory tract anatomy, viral receptor distribution, and clinical disease manifestations to those of humans ^20^. Historically, airborne transmission of H5N1 viruses had not been demonstrated in ferrets prior to the emergence of clade 2.3.4.4b viruses ^17^, thereby intensifying concerns regarding the zoonotic and pandemic potential of currently circulating strains. Notably, human-derived B3.13 genotype H5N1 viruses isolated between 2024 and 2025 were shown to have low-to-efficient transmissibility between ferrets via respiratory droplets ^12,13,17,21^. In contrast, studies on D1.1 genotype H5N1 viruses indicate limited or variable transmissibility in ferrets, with human D1.1 isolates showing no airborne transmission in some studies and variable transmission efficiency following direct contact exposure in others ^17,18^. Thus, although respiratory droplet transmission of B3.13 and D1.1 H5N1 viruses has been investigated, direct contact transmission outcomes appear to vary depending on the genetic composition of the virus. This represents a critical gap in our understanding of mammalian adaptation, transmissibility, and public health risk associated with these emerging HPAIV H5N1 viruses.

In this study, we compared the first human isolates of the emerging and predominant H5N1 clade 2.3.4.4b genotypes B3.13 (A/Texas/37/2024; HPhTX B3.13) and D1.1 (A/Louisiana/12/2024; HPhLA D1.1) in the USA. *In vitro*, we assessed their relative receptor-binding properties, neuraminidase (NA) activities, NA thermostability, and cross-reactivity with reference ferret antisera raised against H5N1 clade 2.3.4.4b candidate vaccine viruses (CVVs). *In vivo*, we compared the infection, pathogenicity and direct contact transmission of both strains in a relevant mammalian ferret model. We identified potential genetic determinants associated with the pathogenicity of the HPhTX B3.13 strain and adaptive mutations acquired by the HPhLA D1.1 strain that may increase its infectivity and transmissibility in ferrets.

## Materials and methods

### Biosafety

All experiments involving HPAIV H5N1 were performed in appropriate biosafety level 3 (BSL3) and animal BSL3 (ABSL3) laboratories at Texas Biomedical Research Institute (Texas Biomed). Animal infection studies were approved by the Texas Biomed Institutional Biosafety (IBC #21-017) and Animal Care and Use (IACUC #1785 MU) committees.

### Cells

Human embryonic kidney (HEK293T), human lung adenocarcinoma epithelial (A549), baby hamster kidney (BHK-21), Crandell-Rees feline kidney (CRFK), Madin-Darby bovine kidney (MDBK), Madin-Darby canine kidney (MDCK), and chicken fibroblast (DF-1) cells were maintained in Dulbecco’s modified Eagle medium (DMEM) (Invitrogen, USA) supplemented with 10% fetal bovine serum (FBS) and 1% PSG (penicillin, 100 U/mL; streptomycin 100 μg/mL; L-glutamine, 2 mM) at 37°C in a humidified 5% CO2 incubator.

### Sera and monoclonal antibodies

The ferret antisera produced against 2.3.4.4b candidate vaccine viruses (CVVs) influenza A/Astrakhan/3212/2020-like (H5N8, clade 2.3.4.4b; IDCDC-RG71A), A/American Wigeon/South Carolina/22-000345-001/2021 (H5N1, clade 2.3.4.4b; IDCDC-RG78A) and A/chicken/Ghana/AVL-763_21VIR7050-39/2021 (H5N1, clade 2.3.4.4b; IDCDC-RG80A), were kindly provided by Drs. Han Di and Bin Zhou at the Center for Disease Control and Prevention (CDC), Atlanta, Georgia, USA.

Monoclonal antibodies (MAbs) raised in mice to detect HPAIV H5N1 were obtained from BEI resources. For Western blot assays, 170-3C12 was utilized to detect PB2; F5-46 was used to detect PB1; and 1F6 was utilized to detect PA. A mouse MAb (HT103; Kerafast, USA) was used to detect viral NP. For immunohistochemical staining, a polyclonal rabbit antibody against the viral NP (PA5-32242; Thermo Fisher Scientific, USA) was used for virus detection in murine lung and brain tissues. A mouse mAb (AC-15; Sigma, USA) was used to detect β-actin and used as a loading control for Western blot.

### Cloning and rescue of recombinant viruses

The eight viral segments of B3.13 influenza A/Texas/37/2024 H5N1 (HPhTX B3.13) were previously described ^11^. Viral segments to generate recombinant D1.1 influenza A/Louisiana/12/2024 H5N1 (HPhLA D1.1) were synthesized in the genetic background of pUC57 (GenScript, USA). The synthesized HPhLA D1.1 sequences were designed according to the published GSAID sequences for the clinical HPhLA D1.1 isolate with Global Initiative on Sharing All Influenza Data (GISAID) accession number EPI3741708-15. The noncoding regions (NCRs) were selected based on assembled sequences for each genomic segment of the HPAIV H5N1 clade 2.3.4.4b viruses to identify the most highly conserved regions among the circulating H5N1 strains. The synthesized viral segments were subcloned and inserted into linearized ambisense pHW2000 to generate pHW-hLAPB2, pHW-hLAPB1, pHW-hLAPA, pHW-hLAHA, pHW-hLANP, pHW-hLANA, pHW-hLAM and pHW-hLANS plasmids ^22,23^. To generate recombinant HPhTX B3.13 or HPhLA D1.1, a complete set of the eight pHW2000 plasmids (1 μg each) was transfected into a coculture of HEK293T/MDCK cells (ratio 3:1) using Lipofectamine™ 3000 Transfection Reagent (Thermo Fisher Scientific, USA) according to the manufacturer’s instructions. Following transfection, the media was replaced with 2 mL of DMEM containing 0.3% bovine serum albumin (BSA), and the cells were incubated in a humidified 5% CO_2_ incubator for 12 h.

In parallel, recombinant PR8-based H5N1 viruses were also generated using the eight-plasmid reverse genetics system as previously described ^24,25^. The six internal gene segments (PB2, PB1, PA, NP, M, and NS) were derived from influenza A/Puerto Rico/8/1934 H1N1 (PR8), whereas the HA and NA viral segments were obtained from either HPhLA D1.1 or HPhTX B3.13. The HA segments were engineered to contain a monobasic cleavage site ^25^. For virus rescue, HEK293T and MDCK cells were co-transfected with 1 μg of each of the eight bidirectional reverse genetics plasmids using the manufacturer’s recommended transfection protocol. At 6 h post-transfection (hpt), the transfection medium was replaced with infection medium consisting of Dulbecco’s modified Eagle medium (DMEM) (Invitrogen, USA) supplemented with 0.3% BSA, 1% penicillin-streptomycin-glutamine (PSG), and 1 μg/mL TPCK-treated trypsin (Sigma-Aldrich, USA). Cultures were maintained at 37°C in a humidified incubator with 5% CO_2_.

At 72 hpt, cell culture supernatants (CCSs) were harvested and centrifuged at 2,500 rpm for 5 min at 4°C. A portion of the collected CCSs was subsequently used to infect 8-10-day-old chicken embryonated Specific Pathogen-Free (SPF) eggs (Charles River Laboratory, USA). At 24h post-inoculation with HPhLA D1.1 or HPhTX B3.13 and 72h post-inoculation with the PR8-based counterparts expressing monobasic HA, allantoic fluids were collected, titrated by the HA assay and the rescued recombinant viruses were aliquoted and stored at −80°C until use.

All viruses were confirmed by whole-genome sequencing of the viral stocks using next-generation sequencing (NextSeq 1000/2000 Illumina). Viral RNA was extracted using the QiAamp Viral RNA Mini Kit (Qiagen, Germany). Sample libraries were prepared using the Illumina Stranded mRNA Prep, Ligation (96 samples) (Illumina, USA) and ran on the NextSeq 1000/2000 P2 XLEAP-SBS Reagent Kit (300 cycles) (Illumina, USA) per the manufacturer’s instructions. All viruses were compared with the published clinical isolates.

### Viral replication

Monolayers of A549, BHK-21, CRFK, MDBK, MDCK, and DF-1 cells were cultured in 6-well plates (∼1x10^6^ cells per well, triplicate) for 24 h, infected with the indicated viruses at a multiplicity of infection (MOI) of 0.001 and kept at 37°C in a humidified 5% CO2 incubator to allow viral adsorption for 1 h. Following viral adsorption, the virus inoculum was removed, and the infected cell monolayers were washed three times with 1XPBS to remove residual non-adsorbed viral particles, and the cell monolayers were then supplemented with 3 mL of infection medium. Infected plates were incubated at 37°C in a humidified 5% CO_2_ incubator. CCSs aliquots (200 µL) were collected at 12, 24, 48, and 72 h post-infection (hpi) and supplemented with an equal volume of fresh infection medium. Viral titers in collected CCSs were determined by plaque assay and immunostaining in MDCK cells as previously described ^26^.

### Neuraminidase (NA) activity and thermostability analysis

NA activity was measured using the MUNANA-based NA-Fluor™ Influenza Neuraminidase Assay Kit (Applied Biosystems, USA) and an enzyme-linked lectin assay (ELLA) as previously described ^27,28^. Viruses were standardized to 32 hemagglutination units (HAU), and serial dilutions were plotted against relative fluorescence units (RFU) for MUNANA or optical density (OD) for ELLA after background subtraction. Each assay was performed in duplicate and repeated three times independently. For thermostability analysis, viruses were standardized to 32 HA units, aliquoted in 100 µL duplicates, and incubated at the indicated temperatures for exactly 10 min using a Bio-Rad CFX96 thermocycler. Residual NA activity was subsequently measured, and each experiment was independently repeated twice in 96-well plates.

### Receptor binding microarrays

A subset of glycans (n = 55) was used as previously described ^28,29^. Recombinant influenza A/Puerto Rico/8/1934(H1N1, PR8) viruses expressing the monobasic HA and NA proteins from HPhLA D1.1 or HPhTX B3.13 were used to conduct the receptor binding microarrays. Viruses were incubated on the arrays and detected using CR6261, a monoclonal anti-HA stalk antibody, followed by an Alexa Fluor™ 647–labeled anti-human secondary antibody. Slides were washed, dried, and scanned, and six technical replicates were analyzed by discarding the highest and lowest values and calculating the mean ± SD of the remaining four replicates.

### Plaque assays and immunostaining

MDCK cells were cultured at a density of ∼1x10^6^ cells per well in 6-well plates and kept overnight at 37°C in a humidified 5% CO_2_ incubator. The following day, cells were infected with 10-fold serial dilutions of the virus/sample for 1 h at 37°C. Following adsorption, cell monolayers were overlaid with post-infection media containing agar and incubated at 37°C in a humidified 5% CO_2_ incubator. At 24-72 hpi, the cells were fixed overnight with 10% neutral buffered formalin solution. For staining and visualization using crystal violet, 1 mL of 1% crystal violet solution was added to each well for 5 min at room temperature (RT) and then rinsed with tap water. For immunostaining, cells were first permeabilized with 0.5% (v/v) Triton X-100 in 1XPBS for 15 min at RT and immunostained with the influenza virus anti-NP mouse MAb HT103 (1:100) and the Vectastain ABC Kit (Vector Laboratories, USA), according to the manufacturer’s instructions. After immunostaining, the plates were scanned and photographed using a ChemiDoc MP Imaging System.

### Microneutralization assays

MDCK cells were seeded in 96-well plates (∼3.5x10^4^ cells/well, quadruplicate) and incubated overnight at 37°C in a humidified 5% CO_2_ incubator. Ferret serum was heat-inactivated prior to use and initially diluted 1:10 in infection medium. Two-fold serial dilutions of ferret serum were then prepared in 96-well plates containing infection medium. An equal volume of infection media containing a total of 100 PFU of virus was added to each serum dilution, and the virus-serum mixtures were incubated for 1 h at RT to allow antibody neutralization. Subsequently, 100 µL of each preincubated virus-serum mixture was added to confluent MDCK cells in 96-well plates. Following incubation for 3 days at 37°C in a humidified 5% CO_2_ incubator, when a cytopathic effect (CPE) was evident, the plates were fixed with 10% neutral buffered formalin solution overnight and stained with 1% crystal violet solution. Neutralizing antibody titers were defined as the reciprocal of the highest serum dilution that completely inhibited virus-induced CPE in MDCK cells.

### Next-generation sequencing

Sequence data were analyzed using a Snakemake v7.32 ^30^ workflow with per-rule Conda environments. All analyses were run on a single compute node and analyses were limited to a maximum of 96 CPU cores and 256 MB of memory limit. Paired-end, raw FASTQ files were quality filtered with Trimmomatic v0.39 ^31^ using the options LEADING:10, TRAILING:10, SLIDINGWINDOW:4:15, and MINLEN:36 to remove low-quality base calls and short reads. Orphaned reads were combined into a file of singleton reads to complement the filtered paired reads. Taxonomic assignment was performed with Kraken2 v2.1.2 ^32^ using the “k2_standard” database and a confidence threshold of 0.2. The database and relevant indices were downloaded from the 15 October 2025 release of https://benlangmead.github.io/aws-indexes/k2 (last accessed 4 March 2026). The reference genomes were indexed with Bowtie2 ^33^ and SAMtools v1.4 ^34^ faidx. Paired and singleton reads were mapped with Bowtie2 using the ‘--very-sensitive-local’ setting. Mapping rates were summarized with SAMtools flagstat, and the depth of coverage was estimated with mosdepth v0.3.2 ^35^. Indel qualities were added to the mapped BAM files using LoFreq v.2.1.5 ^36^ indelqual --dindel, and variants were called per sample with LoFreq using --call-indels, --max-depth 10000, and --no-default-filter. The raw VCFs were filtered to retain variants with a minimum read coverage ≥100, minimum base quality ≥20, and allele frequency ≥0.03. Consensus FASTA sequences were generated from variants with an allele frequency >0.5 using the BCFtools v1.9 ^37^ consensus.

### Minigenome assays

To evaluate viral polymerase activities of HPhTX B3.13, HPhLA D1.1 or related variants *in vitro*, minigenome (MG) assays were performed as previously described ^38^. Briefly, HEK293T cells (12-well plate format, ∼4x10^5^ cells/well, triplicates) were co-transfected using Lipofectamine™ 3000 Transfection Reagent (Thermo Fisher Scientific, USA) with 1 µg of the indicated expression plasmids encoding the viral polymerase proteins PB2, PB1, and PA; and NP. A MG plasmid encoding an influenza viral RNA (vRNA)-like segment expressing ZsGreen (ZsG) fused to nanoluciferase (Nluc) flanked by the NP segment NCR ^38^, was also co-transfected (1 µg). Additionally, a Cypridina luciferase (Cluc)-expressing plasmid (1 µg) was included in all experiments to normalize transfection efficiencies. HEK293T cells transfected with all plasmids except PB1 were used as a negative control. At 6 hpt, the media was replaced with DMEM containing 1% PSG and 10% FBS. Nluc and Cluc expression levels were determined using the Nano-Glo® luciferase assay system (Promega, USA) and the Cypridina luciferase glow assay kit (Thermo Fisher Scientific, USA), respectively, at 30 hpt. Polymerase activity was expressed as a fold change relative to a negative control group lacking PB1, as previously described ^38^. At 30 hpt, live-cell images showing representative ZsG expression were acquired by fluorescence microscopy (EVOS).

### Western blots

HEK293T cell monolayers were washed with 1XPBS and treated with ice-cold NP40 lysis buffer supplemented with a protease inhibitor cocktail (Thermo Fisher Scientific, USA). The cells were lysed on ice and transferred to a microfuge tube. After 30 min, the cells were centrifuged at 15,000 rpm for 15 min at 4°C. The supernatants were mixed with lithium dodecyl sulfate (LDS) sample buffer containing 20% β-mercaptoethanol and heated at 98°C for 5 min before being loaded onto 10% SDS‒PAGE gels. After electrophoresis, the proteins were transferred to nitrocellulose membranes. Next, the membranes were incubated with 5% non-fat dry milk powder in 1XPBS supplemented with 0.05% Tween 20 (PBST) (Sigma, USA). After 1 h of blocking, the membranes were probed with primary antibodies overnight at 4°C. Next, the membranes were washed three times for 5 min each with 5 mL of PBST. Thereafter, the membranes were incubated with secondary antibodies for 2 h at RT. After three additional washes for 5 min each with 5 mL PBST, an enhanced chemiluminescence (ECL) substrate kit was used to visualize chemiluminescence-generated protein bands using SuperSignal (Thermo Fisher Scientific, USA) following the manufacturer’s recommendations.

### Ferret infection and contact transmission study

Outbred 6-month-old castrated male Fitch ferrets were purchased from Triple F Farms (Gillett, USA). Contact transmission studies were performed using four infected ferrets and four naïve contact ferrets for each experimental group. Each cage housed two ferrets: one directly infected ferret and one contact ferret. Briefly, on day 0, the ferrets were anesthetized and intranasally inoculated with 10^4^ PFU of either HPhTX B3.13 (n = 4) or HPhLA D1.1 (n = 4). Infected ferrets were initially housed individually. At 24 hpi (day 1), one naïve ferret was introduced into each cage for direct-contact transmission, resulting in two ferrets per cage. Body weight and temperature were monitored daily throughout the study. Nasal wash samples were collected from anesthetized ferrets every other day for up to 14 days post-infection (DPI) or until animals reached humane endpoint criteria. Serum samples were collected from contact ferrets at 14 DPI or at the time of euthanasia and subsequently subjected to microneutralization assays. Tissue samples, including brain, trachea, lung, heart, kidney, and spleen, were collected at necropsy to determine viral titers via plaque assay. Surviving contact and control ferrets were euthanized at 14 DPI if the humane endpoint criteria had not been reached earlier. Clinical signs were monitored daily and scored using a standardized clinical scoring system. The evaluated parameters included weight loss, upper respiratory signs, dyspnea, mucous membrane appearance, general appearance, activity level, and neurological status/coordination.

Weight loss was scored as follows: 0, 0–2%; 3, >2–10%; 6, >10–20%; and 12, >20%. Upper respiratory signs were scored as 0, normal; 2, sneezing or coughing; 4, ocular and/or nasal discharge; and 8, closed eyes. Dyspnea was scored as 0, normal respiration; 3, increased respiratory rate; 6, mild or transient increased respiratory effort; and 12, marked or persistent respiratory distress or agonal breathing. The presence of a mucous membrane was scored as 0, pink with capillary refill time (CRT) <2 seconds; 6, pale mucous membranes with CRT >2 seconds; and 10, dusky or cyanotic mucous membranes. General appearance was scored as 0, normal; 2, rough hair coat; and 6, rough hair coat with hunched posture. Activity level was scored as 0, active; 2, quiet but responsive to stimulation; 8, moderate lethargy; and 12, moribund or prostrate. Neurological system/coordination was scored as 0, normal; 6, abnormal gait; and 12, seizures or other central nervous system manifestations. Animals with a cumulative clinical score of ≥6 demand Alert Clinical Vet for increased monitoring and clinical intervention. Animals that reached a cumulative clinical score of ≥12 were humanely euthanized according to institutional animal care and use guidelines.

### Histopathology and immunohistochemistry

Tissues from euthanized ferrets were fixed in 10% neutral buffered formalin for a minimum of 24 h, transferred to 75% ethanol, paraffin-embedded, sectioned (4 µm), and stained with hematoxylin and eosin (H&E) as previously described ^38^. Stained sections were evaluated by a board-certified veterinary pathologist in a blinded manner. Whole-slide images were acquired at 20X magnification (Axio Scan Z1, Zeiss) and analyzed using HALO software (Indica Labs, USA) to quantify pathology. The tissues were counterstained using Hematoxylin (Roche, USA) followed by Bluing Reagent (Roche, USA). Immunohistochemistry was performed on formalin-fixed tissue sections to detect viral NP antigen in tissues by incubating with an influenza NP rabbit polyclonal antibody (Invitrogen, USA) at a concentration of 1:1,500 for 1 h at RT and immunostaining using anti-rabbit HQ (Roche, USA) for 8 min, followed by incubation with anti-HQ HRP (Roche, USA) for 8 min at 36°C. Influenza A NP was then visualized using ChromoMAP DAB (Roche, USA). Total cell counts positive for immunostaining were normalized to the tissue area and analyzed using HALO software (Indica Labs, USA).

### Computational docking methodology

Systems were prepared from crystal structures (PDB: 8RMR, 8R1L) ^39,40^ with mutations introduced via CHARMM-GUI ^41,42^. All-atom molecular dynamics simulations were conducted using GROMACS 2025.2 ^43^ with the CHARMM36m force field ^44^. The systems were equilibrated (energy minimization, 100 ps NVT, 100 ps NPT at 310 K) and simulated for 100 ns in the NPT ensemble. Binding free energies were calculated using gmx_MMPBSA ^45^ over 200 trajectory snapshots (frames 1-1000, 5-frame intervals). RMSD, RMSF, and contact analysis (4.5 Å cutoff) were performed using MDAnalysis ^46–48^. Epistatic interactions were quantified as deviations from additivity with significance determined by bootstrap resampling of trajectory frames. Because MM/PBSA is an end-state approximation, binding energies are interpreted comparatively rather than as absolute affinities.

## Results

### *In vitro* characterization and comparison of clade 2.3.4.4b genotype B3.13 and D1.1 H5N1 viruses

We evaluated the replication efficiency *in vitro* and the pathogenicity and transmissibility of the human-derived clade 2.3.4.4b genotype B3.13 A/Texas/37/2024 H5N1 (HPhTX B3.13) with clade 2.3.4.4b genotype D1.1 A/Louisiana/12/2024 H5N1 (HPhLA D1.1) (**Fig. 1**). The B3.13 genotype was initially transmitted from migratory birds to cattle and subsequently to humans ^1,2^. During circulation at the cattle–human interface, some viruses acquired mammalian-adaptive mutations, including the PB2 627K mutation ^1,24^. In contrast, the D1.1 genotype primarily circulated from migratory birds to poultry and then to humans ^1,49^. More recently, however, D1.1 genotype has also been reported to transmit from poultry to cattle and subsequently to humans ^1,49^ (**Fig. 1A**). Interestingly, recombinant HPhTX B3.13 and HPhLA D1.1 exhibited distinct plaque morphologies and sizes on permissive MDCK cell monolayers (**Fig. 1B**). HPhTX B3.13 produced larger, well-defined plaques with clear boundaries, consistent with more efficient cell-to-cell spread and/or higher cell toxicity *in vitro* (**Fig. 1C**). In contrast, HPhLA D1.1 formed smaller plaques with less sharply demarcated edges, suggesting relatively reduced spread kinetics under the same experimental conditions (**Fig. 1C**). Consistently, HPhTX B3.13 replicates to significantly higher titers in mammalian (A549, BHK21, CRFK, MDBK, and MDCK) but not avian DF1 cells (**Fig. 1C**). These findings indicate a clear replication advantage of HPhTX B3.13 over HPhLA D1.1 in mammalian cells, while replication in avian cells was not significantly different under the same experimental conditions for both H5N1 viruses.

**Figure 1.**
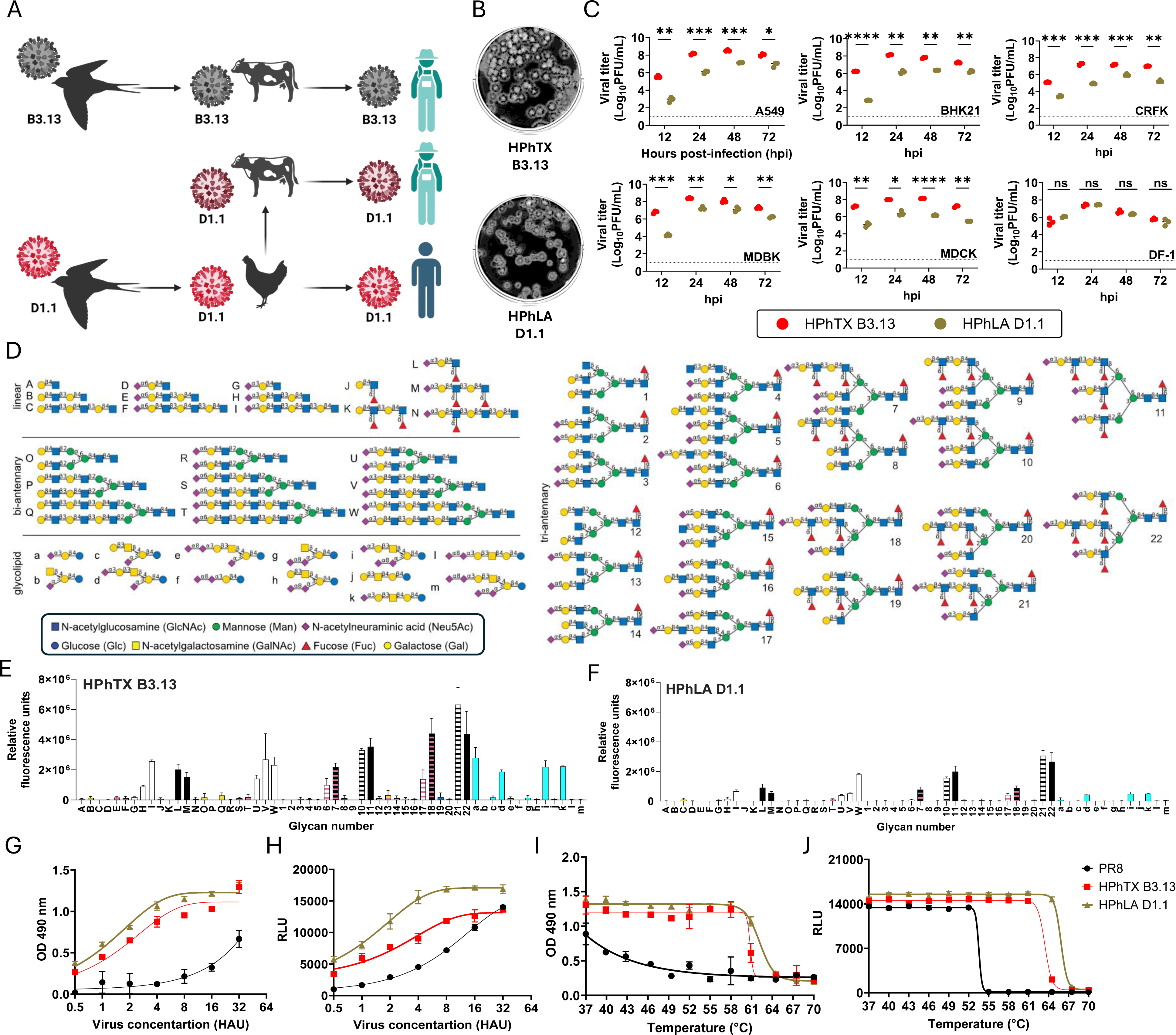
*In vitro* characterization of clade 2.3.4.4b genotype B3.13 and D1.1 H5N1 viruses. **(A) Schematic of the inferred natural transmission pathways:** H5N1 genotype B3.13 was initially transmitted from migratory birds to cattle and subsequently to humans, with some viruses acquiring mammalian-adaptive mutations during circulation at the cattle–human interface. H5N1 genotype D1.1 predominantly circulated from migratory birds to poultry and then to humans; more recently, transmission from poultry to cattle and subsequently to humans has also been reported. **(B) Plaque phenotypes of HPhTX B3.13 and HPhLA D1.1. (C) Replication kinetics:** Indicated mammalian and avian cells were infected (MOI 0.001) with HPhTX B3.13 or HPhLA D1.1 and at 12, 24, 48, and 72 hpi, CCSs were collected to evaluate the presence of the virus by plaque assay. **(D-F) Glycan microarray analysis of receptor-binding specificity for HPhTX B3.13 and HPhLA D1.1 viruses:** Representative glycan structures included in the microarray (**D**). Binding profiles of HPhTX B3.13 (**E**) and HPhLA D1.1 (**F**). Fluorescence intensity corresponds to virus binding to the indicated glycan structures. Key: α2,3-SA, α2-3-sialyl-N-acetyllactosamine (avian-type receptor); α2,6-SA, α2-6-sialyl-N-acetyllactosamine (human-type receptor). **(G-H) NA activity of HPhTX B3.13 and HPhLA D1.1 measured using (G) MUNANA and (H) enzyme-linked lectin assay (ELLA).** Virus preparations were normalized to 32 hemagglutination units (HAU) prior to analysis. Data are presented as log10-transformed relative fluorescence units (RFU) for MUNANA or absorbance at 490 nm for ELLA and are shown as mean ± SD. Experiments were performed independently twice, each in duplicate. **(I-J) NA thermostability of HPhTX B3.13 and HPhLA D1.1 assessed using (I) MUNANA and (J) ELLA:** Virus preparations were normalized to 32 HAU before heat treatment. Residual NA activity is presented as mean ± SD from two independent experiments performed in duplicate. HPhTX B3.13: A/Texas/37/2024 H5N1. HPhLA D1.1: A/Louisiana/12/2024 H5N1, PR8: A/Puerto Rico/1/1934 H1N1.

### Differential replication in mammalian cells is not associated with altered receptor binding specificity or enhanced NA function

To investigate the potential mechanisms underlying the observed differences in mammalian replication, we assessed HA receptor-binding specificity using glycan microarray analysis (**Figs. 1D-F**). No significant differences in receptor-binding profiles were detected between HPhTX B3.13 and HPhLA D1.1. Both viruses retained a strong preference for avian-type sialic acid receptors and showed minimal binding to human-type receptors, indicating that altered receptor specificity does not account for the enhanced replication of HPhTX B3.13. We next evaluated NA activity and thermostability using ELLA and MUNANA assays (**Figs. 1G-J**). Surprisingly, HPhTX B3.13 exhibited slightly lower NA activity than HPhLA D1.1 across both assay platforms (**Figs. 1G-H**). Despite this difference in enzymatic activity, the NA proteins of both viruses remained highly stable across a broad temperature range (37-58°C) and lost activity only at temperatures exceeding 67°C (**Figs. 1I-J**). Notably, greater NA thermostability for HPhLA D1.1 at 61°C was observed by MUNANA and ELLA and at 64°C by MUNANA, suggesting some degree of substrate-dependent variation in enzyme stability. Taken together, these results indicate that the enhanced replication and larger plaque phenotype of HPhTX B3.13 are not associated with changes in receptor-binding specificity or increased NA activity. Instead, the lower NA activity observed in HPhTX B3.13 may contribute to improved virion retention at the cell surface, thereby facilitating more efficient cell-to-cell spread and plaque expansion *in vitro*.

### Antigenic relatedness of clade 2.3.4.4b genotypes B3.13 and D1.1 to the current CDC candidate vaccine virus (CVV)

Cross-reactivity analyses using ferret antisera raised against CDC CVV strains demonstrated that both HPhTX B3.13 and HPhLA D1.1 H5N1 viruses are antigenically related to the corresponding CVVs. Microneutralization (MN) assays revealed comparable neutralizing capacities of the ferret sera against both viruses, indicating no substantial antigenic differences between HPhTX B3.13 and HPhLA D1.1, at least in this assay (**Table 1**), suggesting the feasibility of using the three current CDC CVVs (IDCDC-RG71A, -RG78A, and -RG80A) to provide protection against the B3.13 and D1.1 H5N1 genotypes.

**Table 1.** Cross reactivity with ferret positive sera raised against CDC CVV strains.

| Sera | MN titer against influenza<br>A/Texas/37/2024<br>H5N1 B3.13 |  |  | MN titer against Influenza<br>A/Louisiana/12/2024<br>H5N1 D1.1 |  |  |
| --- | --- | --- | --- | --- | --- | --- |
| Ctrl ferret sera | <10 | <10 | <10 | <10 | <10 | <10 |
| IDCDC-RG71A* | 10240 | 10240 | 10240 | 10240 | 10240 | 10240 |
| IDCDC-RG78A** | 2560 | 2560 | 2560 | 2560 | 2560 | 2560 |
| IDCDC-RG80A*** | 20480 | 20480 | 20480 | 20480 | 20480 | 20480 |

\* IDCDC-RG71A: A/Astrakhan/3212/2020 H5N8
\*\* IDCDC-RG78A: A/American Wigeon/South Carolina/22-000345-001/2021 H5N1
\*\*\* IDCDC-RG80A: A/chicken/Ghana/AVL-763\_21VIR7050-39/2021 H5N1

### Pathogenicity and transmission of clade 2.3.4.4b genotype B3.13 and D1.1 H5N1 viruses in ferrets

We next evaluated the virulence and transmissibility of the HPhTX B3.13 and HPhLA D1.1 H5N1 viruses in ferrets (**Fig. 2A**). Infection with HPhTX B3.13 resulted in more pronounced body weight loss than infection with HPhLA D1.1 (**Figs. 2B** & **2D**). This increased pathogenicity was also associated with earlier onset of mortality in both HPhTX B3.13 infected ferrets and their direct-contact counterparts (**Figs. 2C** & **2E**). Furthermore, the transmissibility of HPhTX B3.13 was markedly greater than that of HPhLA D1.1, as indicated by elevated clinical scores (**Fig. 3A**) and sustained fever (**Fig. 3B**) in both infected and contact ferrets. Notably, all contact ferrets in the HPhTX B3.13 group died between 6 and 8 DPI (**Fig. 3C**). In contrast, among HPhLA D1.1 contact ferrets, only one succumbed to infection (**Fig. 3E**), one showed a slight increase in body temperature (F17), and the other two ferrets remained asymptomatic (**Fig. 3E**). These results demonstrate that HPhTX B3.13 is not only more pathogenic but also more efficiently transmitted among ferrets than HPhLA D1.1.

**Figure 2.**
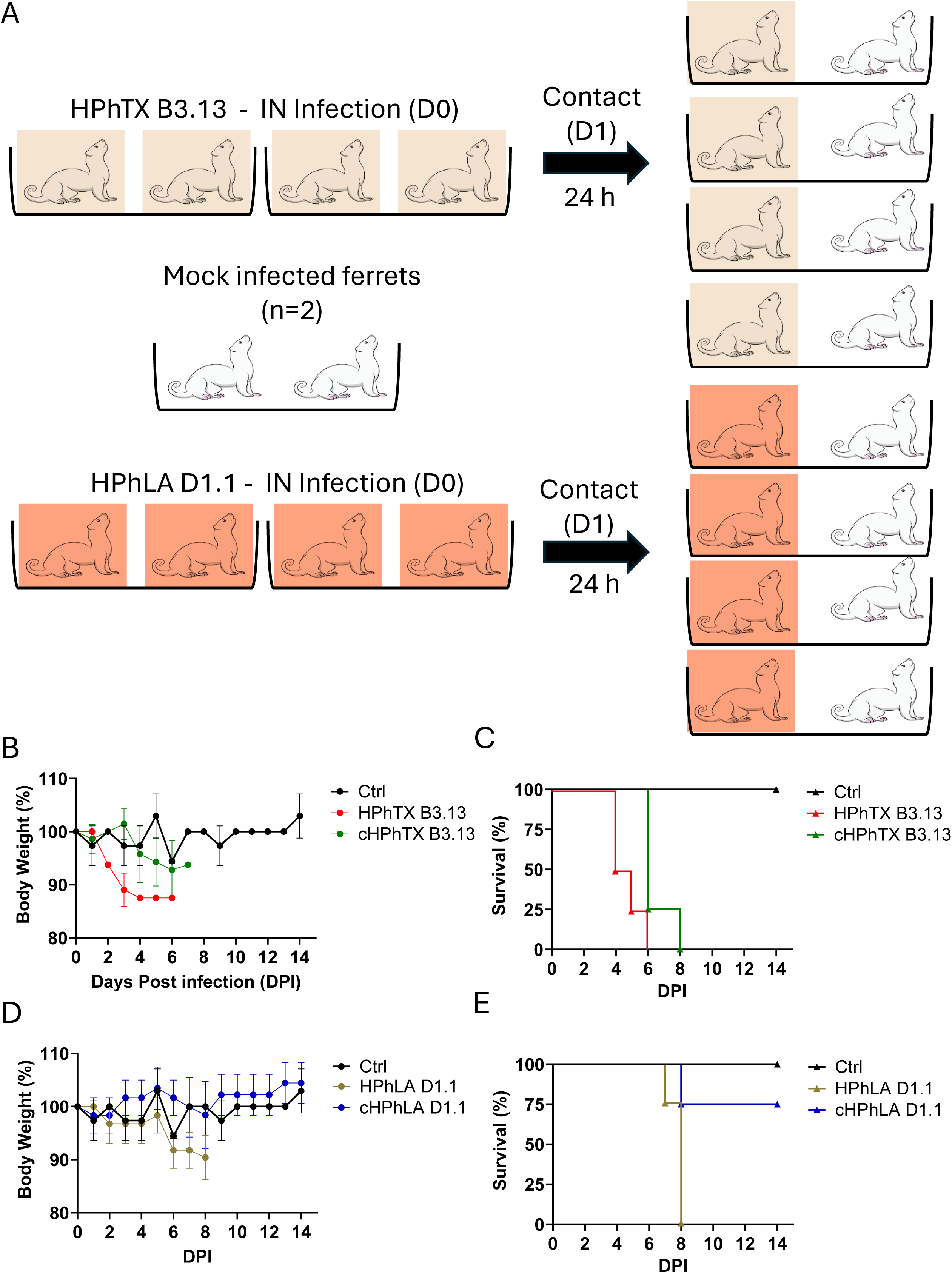
HPhTX B3.13 causes increased virulence and transmissibility in ferrets. **(A) Experimental design for ferret infection and transmission studies:** Ferrets were infected intranasally with 10^4^ PFU of HPhTX B3.13 or HPhLA D1.1, and direct-contact naïve ferrets were introduced in the same cage 24 h after infection. **(B & D) Body weight changes:** Changes in body weight over time in infected and contact ferrets infected with HPhTX B3.13 (**B**) and HPhLA D1.1 (**D**). **(C & E) Animal survival:** survival curves for infected and contact ferrets infected with HPhTX B3.13 **(C)** and HPhLA D1.1 **(E).** Data are presented as mean ± SEM (n = 4). HPhTX B3.13: A/Texas/37/2024 H5N1 infected ferrets. HPhLA D1.1: A/Louisiana/12/2024 H5N1 infected ferrets. cHPhTX B3.13: A/Texas/37/2024 H5N1 contact ferrets. cHPhLA D1.1: A/Louisiana/12/2024 H5N1 contact ferrets. Ferret cage icons are color-coded according to experimental group: yellow indicates ferrets infected with HPhTX B3.13, orange indicates ferrets infected with HPhLA D1.1, and uncolored icons indicate naïve control ferrets or direct-contact ferrets, as specified for each panel.

**Fig 3.**
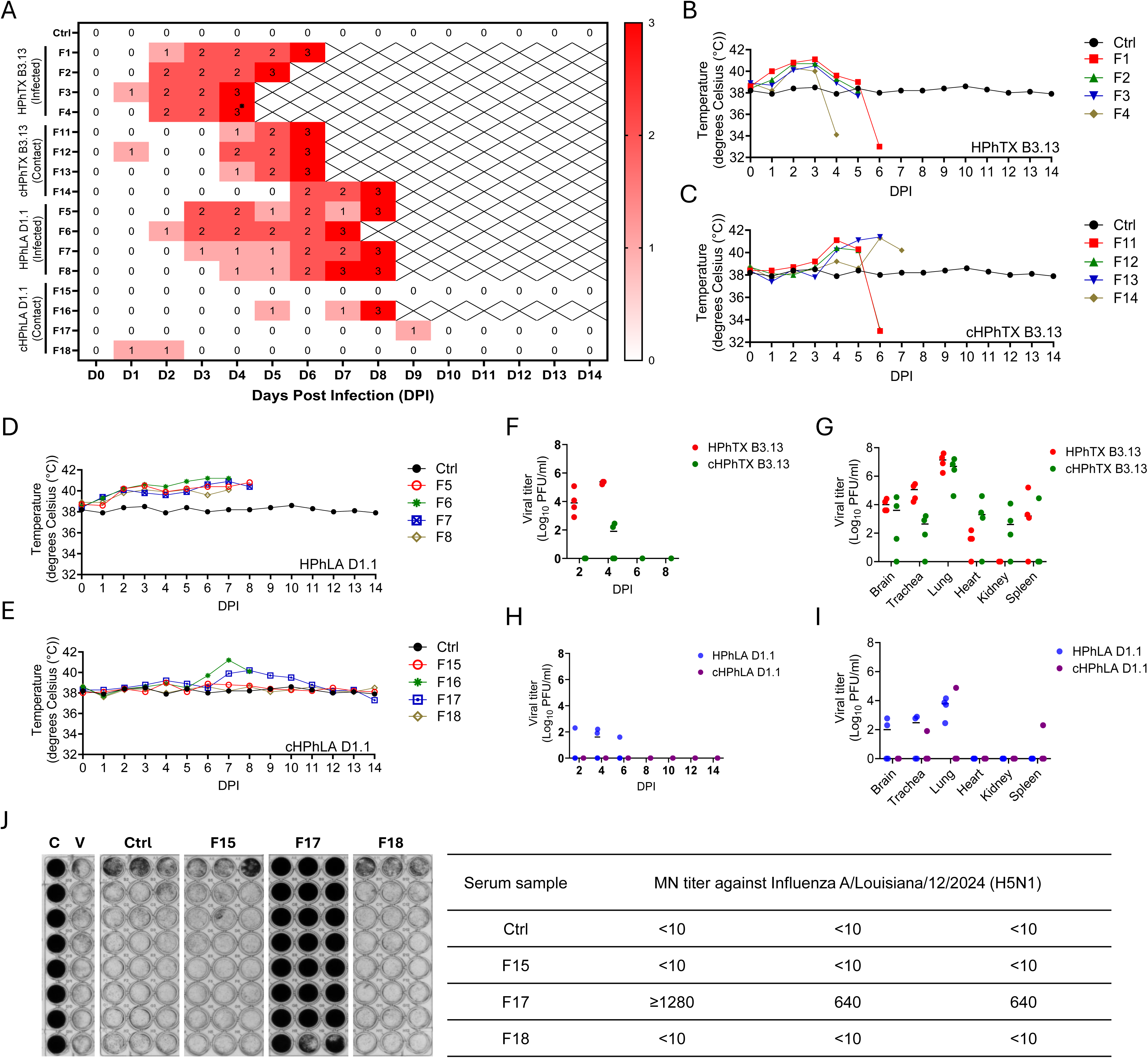
Clinical scores and body temperatures of infected and contact ferrets following infection with HPhTX B3.13 and HPhLA D1.1. **(A) Clinical scores of HPhTX B3.13 and HPhLA D1.1 infected ferrets and their contact animals**: Clinical scores were assigned as follows: 0 = no clinical signs; 1 = mild clinical signs (Clinical score of ≥1 to <6); 2 = moderate clinical signs (alert score; Clinical score of ≥6 to <12); and 3 = humane endpoint requiring euthanasia (Clinical score of ≥12). D, days post-infection (dpi); F, ferret number. **(B-C) Body temperatures of HPhTX B.13 infected (B) and contact (C) ferrets. (D-E) Body temperatures of HPhLA D1.1 infected (D) and contact (E) ferrets. (F-I) Viral shedding and tissue tropism of HPhTX B3.13 and HPhLA D1.1 in infected and contact ferrets:** Nasal washes were collected from infected and contact ferrets at the indicated time points post-infection to assess viral shedding (**F** & **H**). Virus titers in various tissues from HPhTX B3.13 and HPhLA D1.1 infected and contact ferrets, respectively, including lungs, trachea, brain, heart, kidney, and spleen, were determined by plaque assay (**G** & **I**). **(J) Microneutralization (MN) assays:** Microneutralization results of surviving contact ferrets revealed that only one ferret (F17) seroconverted, while the other two remained seronegative. HPhTX B3.13: A/Texas/37/2024 H5N1 infected ferrets. HPhLA D1.1: A/Louisiana/12/2024 H5N1 infected ferrets. cHPhTX B3.13: A/Texas/37/2024 H5N1contact ferrets. cHPhLA D1.1: A/Louisiana/12/2024 H5N1 contact ferrets. * Nasal wash for this ferret was not collected.

### Replication of clade 2.3.4.4b genotype B3.13 and D1.1 H5N1 viruses in ferrets

To investigate viral replication, shedding, and tissue distribution, nasal washes and various organs were collected from infected and direct-contact ferrets at defined times post-infection (**Fig. 3**). HPhTX B3.13 infected ferrets exhibited robust viral shedding in nasal washes, with high virus titers detected as early as 2 DPI (**Fig. 3F**). Analysis of tissue tropism revealed that HPhTX B3.13 replicated efficiently in multiple organs, including the lungs, trachea, brain, heart, kidney, and spleen (**Fig. 3G**). In contrast, HPhLA D1.1-infected ferrets presented lower nasal virus titers, and shedding was largely restricted to early time points (**Fig. 3H**). In addition, viral titers in the lungs, trachea, brain, heart, kidney, and spleen tissues were consistently lower in HPhLA D1.1 infected ferrets than in HPhTX B3.13 infected animals, which exhibited more limited organ dissemination and lower viral loads (**Fig. 3I**). Importantly, direct-contact ferrets exposed to HPhTX B3.13 also exhibited substantial viral replication in both nasal washes and tissues, confirming efficient viral transmission (**Figs. 3F-G**). In contrast, HPhLA D1.1 contact ferrets exhibited minimal or undetectable viral replication, reflecting decreased viral transmissibility (**Figs. 3H-I**). Serological analysis via the MN assay revealed that among the surviving contact ferrets, one animal (F17) infected with HPhLA D1.1 seroconverted, while the other two remained seronegative (**Fig. 3J**). These data indicate that HPhTX B3.13 not only causes higher viral loads and broader tissue dissemination but is also more efficiently transmitted to naïve ferrets than HPhLA D1.1. Collectively, these results demonstrate that differences in viral replication and shedding likely contribute to the observed disparities in virulence and transmissibility between the HPhTX B3.13 and HPhLA D1.1 H5N1 strains.

### Histopathology and viral antigen distribution in infected ferrets

Tracheal, lung, brain, and spleen tissues were collected at necropsy upon reaching predefined clinical endpoints and were examined histologically and by immunohistochemistry (IHC) for influenza viral NP antigen (**Fig. 4**).

**Figure 4.**
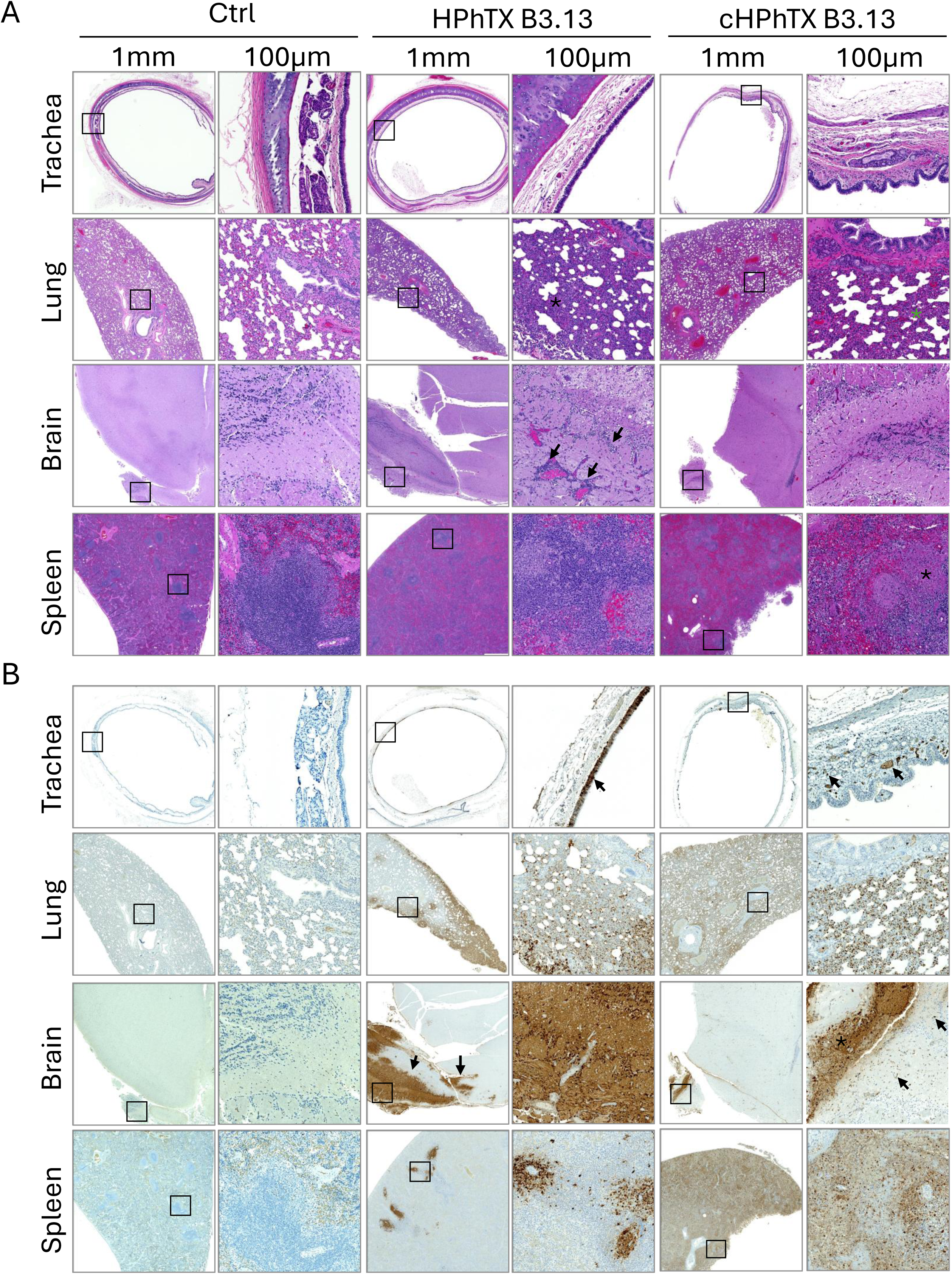
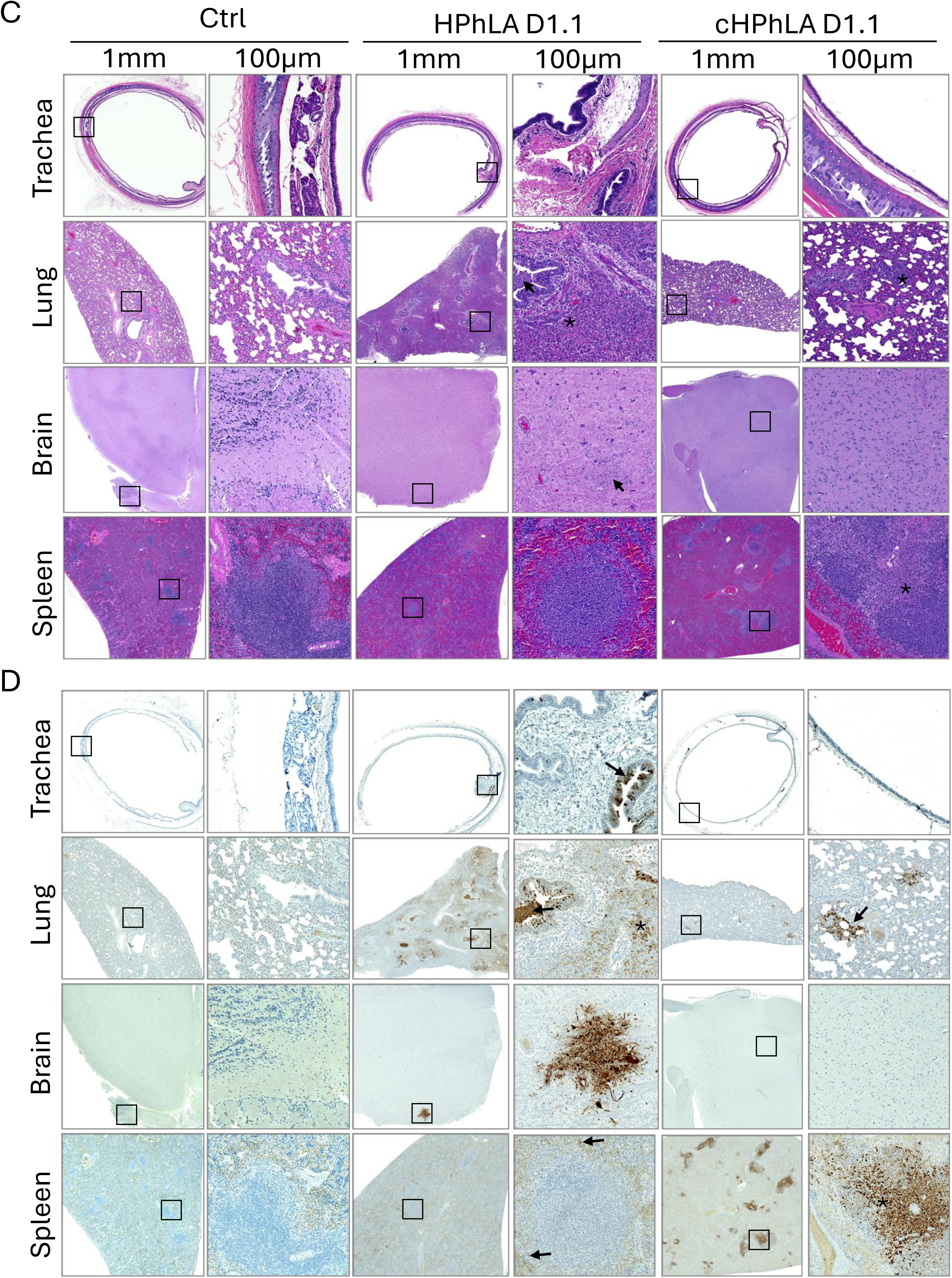
Histopathologic lesions and influenza viral NP antigen distribution in tissues of ferrets infected with HPhTX B3.13 and HPhLA D1.1 H5N1 viruses. Ferrets infected with HPhTX B3.13 or HPhLA D1.1 and naïve contact ferrets were introduced at 24 hpi. Trachea, lung, brain, and spleen tissues were collected for histopathology and IHC detection of viral NP antigen. **(A-D) Histopathologic lesions and NP antigen distribution in HPhTX B3.13 and HPhLA D1.1 infected ferrets:** Viral antigen was detected in respiratory, neural, and lymphoid tissues with more extensive distribution in HPhTX B3.13 infected ferrets (**A** & **B**) compared to HPhLA D1.1 infected ferrets (**C** & **D**). Whole-slide images were acquired using a Zeiss Axioscan Z.1 scanner and analyzed using HALO-AI software. Scale bars: 1 mm (first and third columns) and 100 µm (second and fourth columns). Black arrows indicate representative influenza virus NP antigen-positive cells or specific histopathological lesions. Black and green asterisks denote representative pathological changes or areas of viral antigen staining highlighted in each panel. Squares outline regions that are shown at higher magnification.

Both directly infected and contact HPhTX B3.13 ferrets exhibited mild tracheal submucosal infiltrates of lymphocytes and plasma cells and interstitial pulmonary inflammation (**Fig. 4A**). In the directly infected ferret, segmental lung lesions contained prominent infiltrates of lymphocytes, plasma cells and macrophages along with some fibrin within the alveolar septa (**Fig. 4A, black asterisk)**; the contact ferret showed milder infiltration with diffuse, moderate congestion (**Fig. 4A**). The directly infected ferret exhibited mild perivascular cuffing, gliosis and occasional satellitosis, primarily within the olfactory lobe and frontal cortex, whereas no histopathologic brain lesions were detected in the contact ferret. The spleens of both animals exhibited lytic necrosis of the lymphocytes (lymphocytolysis) within the periarteriolar lymphoid sheath (PALS); however, there was multifocal and mild to moderate infiltration in the directly infected ferret, and the contact ferret showed marked lymphocytolysis throughout the spleen (**Fig. 4A)**. NP antigen was detected in the trachea, lung, brain, and spleen of both ferrets, including epithelial, inflammatory, neural, endothelial, and splenic lymphoid compartments (**Fig. 4B**)

Similarly, both HPhLA D1.1 ferrets (directly infected and in contact) exhibited small infiltrates of lymphocytes and plasma cells throughout the submucosa (**Fig. 4C**). The lungs of the HPhLA D1.1 infected ferret exhibited few bronchioles with attenuated or sloughed epithelium filling the lumen with karryorhectic and cellular debris, infiltrating macrophages, lymphocytes, and plasma cells (**Fig. 4C, Row 2, black arrow**).

Surrounding the affected bronchioles are areas of consolidation with infiltration of abundant lymphocytes, plasma cells, macrophages, and alveolar septa with type II pneumocyte hyperplasia. The lung from the contact infected ferret showed diffuse, mild infiltration of mononuclear inflammatory cells expanding the alveolar septa. The brain from infected ferret showed multiple small areas of gliosis, and infiltration of some lymphocytes, plasma cells and few neutrophils (**Fig. 4C, Row 3, black arrow**). No significant lesions were observed in the brain of the contact infected ferret (**Fig. 4C, Row 3**). The spleen of infected ferret did not show any significant lesions, whereas the contact infected ferret showed multifocal areas of moderate lytic necrosis of the lymphocytes (asterisk) within the periarteriolar lymphoid sheath (PALS) (**Fig. 4C, Row 4**). IHC for the detection of the influenza viral NP antigen revealed the presence of the viral antigen in multiple tissues from infected and contact HPhLA D1.1 ferrets (**Fig. 4D**). The trachea from infected ferret showed scattered viral antigen staining (brown color) in the epithelial cells (black arrow) (**Fig. 4D, Row 1**). The contact infected ferret trachea showed no viral antigen staining in the epithelium and rare staining of cells in the submucosa (**Fig. 4D, Row 1**). Marked viral antigen staining along alveolar septa within areas of consolidation (black asterisk) and including the debris (black arrow) within the bronchi and bronchioles was observed in the lungs of infected ferrets, whereas multifocal, small patches of viral antigen staining of alveolar septa (black arrow) were observed in contact infected ferret (**Fig. 4D, Row 2**). The brain from infected ferret showed mild viral antigen staining in the brainstem while contact infected ferret did not show any viral antigen within the brain (**Fig. 4D, Row 3**). The spleen from infected ferret had mild scattered viral antigen-stained cells throughout (black arrow) but the contact infected ferret had multifocal, moderate viral antigen staining in the PALS areas (black asterisk) (**Fig. 4D, Row 4**). The control tissues showed no inflammatory lesions and no viral NP antigen staining (**Figs. 4A-D**).

Collectively, these results indicate that both H5N1 genotypes can disseminate beyond the respiratory tract following direct and contact infection. However, HPhTX B3.13 was associated with more extensive histopathological changes and greater influenza NP antigen distribution across multiple organs than HPhLA D1.1, particularly after contact transmission, consistent with its greater pathogenicity and transmission efficiency in ferrets.

### Genetic stability of HPhTX B3.13 and HPhLA D1.1 in infected and contact ferrets

We next performed sequence analysis of lung homogenates from ferrets infected with the HPhTX B3.13 and HPhLA D1.1 H5N1 strains. The original NGS data were deposited into BioProject #PRJNA1475507. Compared to the inoculated virus (**Supplementary Fig. 1A**), HPhTX B3.13 did not exhibit evidence of positive selection for specific mutations following replication or transmission and remained highly lethal in both directly infected (**Table 2** & **Supplementary Figs. 1B-E**) and contact (**Table 2** & **Supplementary Figs. 1F-I**) ferrets. The NP G49S substitution was identified in only one of the four contact ferrets and was not detected in the remaining contact or directly infected animals, suggesting that it arose as an isolated event and is unlikely to represent a transmission-associated adaptive mutation. In contrast, HPhLA D1.1 (**Supplementary Fig. 1J**) consistently acquired the well-characterized mammalian adaptation marker PB2 E627K in all directly infected ferrets, even after a single round of infection (**Table 2** & **Supplementary Figs. 1K-N**). Furthermore, HPhLA D1.1 was transmitted to two contact ferrets, although only one of these animals developed severe disease. Similarly, sequencing of lung and spleen homogenates from a contact ferret identified two nonsynonymous substitutions (E627K and Q194K) in the PB2 protein and one synonymous mutation in the M gene (C320T; L99L) (**Table 2** & **Supplementary Figs. 1O** & **1P**). After this sequence analysis, we further investigated the potential contribution of these mutations to viral pathogenicity and transmissibility using plasmid-based minigenome (MG) assays, with particular emphasis on the previously uncharacterized Q194K substitution.

**Table 2.**
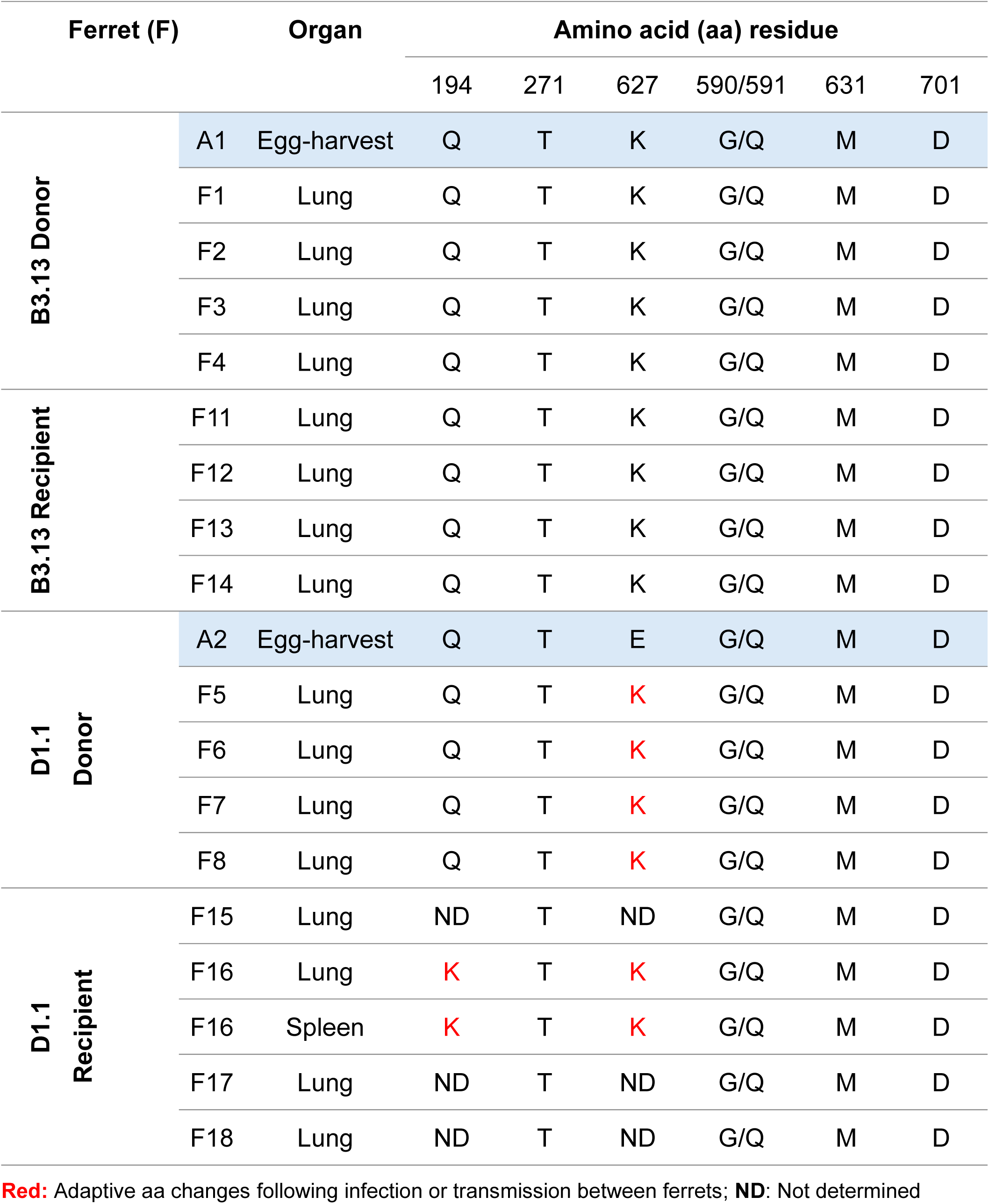
Mammalian adaptation markers and adaptive mutations identified in PB2 segments by NGS of tissue homogenates from infected and contact ferrets.

### PB2 is the major determinant of enhanced polymerase activity in HPhTX B3.13 with PB2 residues 627K and 194K being associated with improved polymerase activity and ferret-to-ferret transmission of HPhLA D1.1

To compare the polymerase activities of the HPhTX B3.13 (TX) and HPhLA D1.1 (LA) H5N1 viruses, we employed a previously described dual-reporter MG assay in HEK293T cells (**Supplementary Fig. 2A**) ^38^. Compared to those of HPhLA D1.1, vRNP and negative control transfections missing the PB1 segments of HPhLA D1.1 (D1.1 LA-PB1) or HPhTX B3.13 (B3.13 TX-PB1), enhanced polymerase activity was detected in HPhTX B3.13 vRNP as quantified by Nluc activity normalized to Cluc (**Fig. 5A**). Consistently, a remarkable increase in ZsG fluorescence was detected with HPhTX B3.13 vRNP (**Supplementary Fig. 2B**). Comparable expression levels of PB2, PB1, PA, and NP were confirmed by Western blot analysis in comparison with the pCAGGS-transfected cell lysate as a negative control (Ctrl) (**Fig. 5B**), indicating that differences in reporter activity were not attributable to differences in protein expression (**Supplementary Table 1**). To identify the viral gene segment responsible for the enhanced polymerase activity of HPhTX B3.13, individual polymerase gene segments were exchanged between HPhTX B3.13 and HPhLA D1.1. Substitution of the HPhLA D1.1 PB2 segment with the corresponding HPhTX B3.13 PB2 significantly increased polymerase activity, identifying HPhLA D1.1 PB2 as a limiting determinant of polymerase function (**Fig. 5C** & **Supplementary Fig. 2C**). Conversely, replacement of the HPhTX B3.13 PB2 segment with that of HPhLA D1.1 reduced polymerase activity, further demonstrating that PB2 is a major determinant of the enhanced polymerase activity associated with the HPhTX B3.13 genotype (**Fig. 5D** & **Supplementary Fig. 2D**). To investigate whether adaptive PB2 mutations contributed to the observed differences, we generated plasmids expressing HPhLA D1.1 PB2 mutants harboring host adaptation-associated substitutions identified in naturally occurring H5N1 isolates (**Fig. 5E**). MG assays demonstrated that the introduction of the mammalian adaptive PB2 mutations 627K and 701N enhanced viral polymerase function compared to that of the HPhLA D1.1 WT PB2 (**Fig. 5F** & **Supplementary Fig. 2E**). Compared to those of the pCAGGS-transfected cell lysate (Ctrl), the western blot analysis confirmed comparable expression of WT and mutant PB2 proteins. These findings further indicate that the observed functional differences were not due to altered protein expression (**Fig. 5G**). To further investigate the impact of ferret adaptive PB2 mutations on viral polymerase activity, the effects of PB2 substitutions 627K, 194K, and the combined 627K/194K mutant were evaluated at 33°C (**Figs. 5H** & **5I**), 37°C (**Figs. 5J** & **5K**), and 39°C (**Figs. 5L** & **5M**) using the same MG assay. Polymerase activity, quantified by Nluc expression and confirmed by ZsG expression, revealed temperature-dependent differences among the PB2 variants. Notably, the PB2 627K single and 627K/194K double mutants exhibited consistently greater polymerase activity than the HPhLA D1.1 WT PB2 at all the temperatures tested, approaching the polymerase activity of HPhTX B3.13. In contrast, the PB2 194K mutation alone significantly enhanced polymerase activity only at 37°C and 39°C, with no significant effect at 33°C. This temperature-dependent increase may help explain the limited viral shedding and reduced transmission efficiency of HPhLA D1.1 compared with HPhTX B3.13, suggesting that PB2 194K alone is insufficient to fully support efficient viral replication in the cooler upper respiratory tract. Taken together, these findings identify PB2 as a major determinant of polymerase activity and demonstrate that mammalian adaptive PB2 mutations, particularly the combined 627K/194K substitutions, substantially enhance the polymerase function of H5N1 D1.1 viruses.

**Fig 5.**
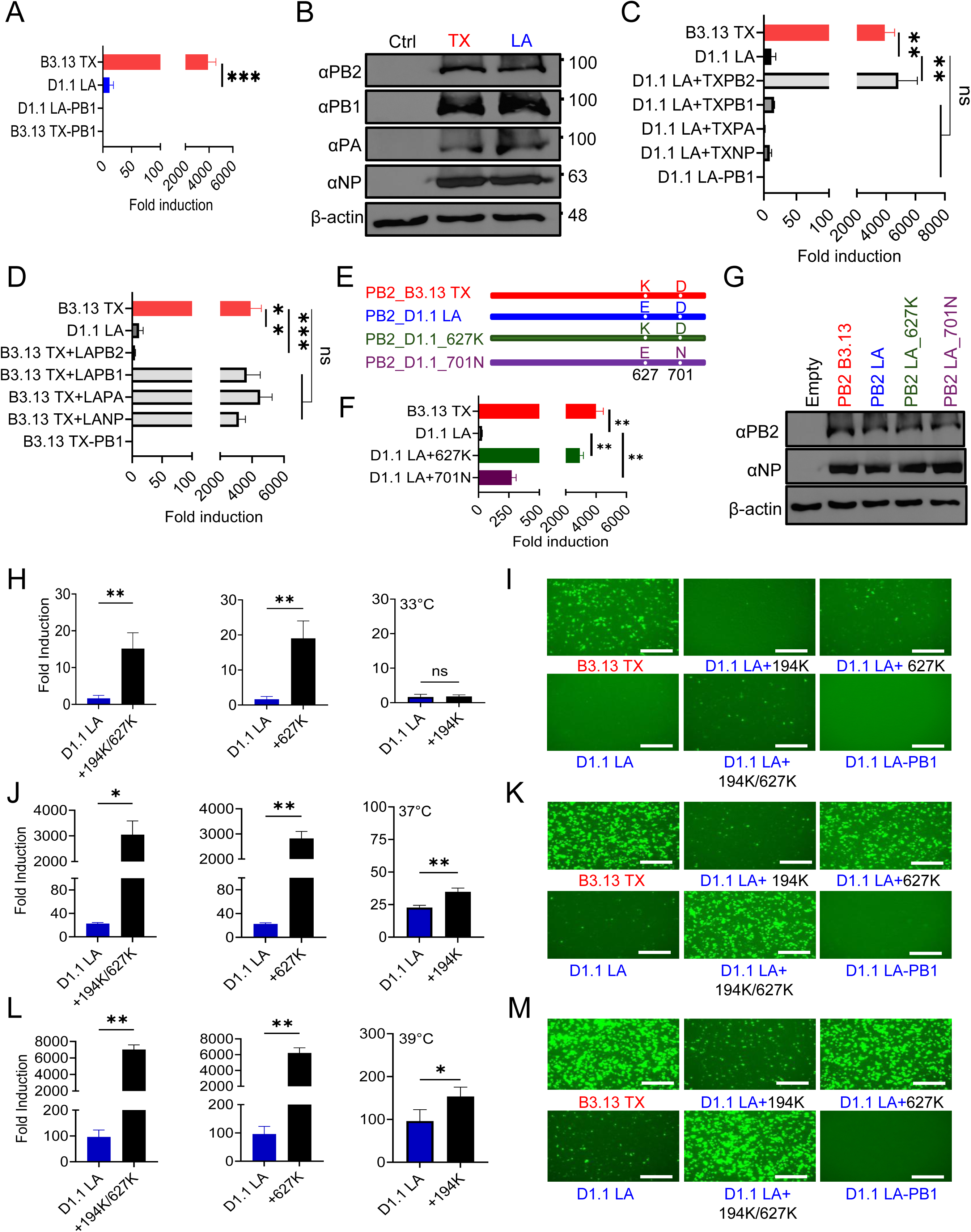
Enhanced polymerase activity of HPhTX B3.13 (TX) and HPhLA D1.1 mutant 627K/194K compared to HPhLA D1.1 WT. (A) Comparison of HPhTX B3.13 and HPhLA D1.1 minigenome activities: HEK293T cell supernatants transfected with the MG, together with the plasmids encoding the viral ribonucleoprotein (vRNP) complex (PB2, PB1, PA, and NP) of HPhTX B3.13 (TX) and HPhLA D1.1 (LA), and a Cluc expression plasmid to normalize transfection efficiency, were collected at 30 hpt to assess viral genome replication and gene transcription. Nluc activity was measured and normalized to Cluc activity and expressed as fold induction relative to a control group lacking the respective PB1 plasmid (–PB1). **(B) Western blot analysis:** HEK293T cell lysates transfected in (A) were used to assess HPhTX B3.13 (TX) and HPhLA D1.1 (LA) PB2, PB1, PA, and NP expression. Empty pCAGGS plasmid-transfected cells were included as negative controls. β-actin was included as a loading control. Molecular markers are indicated on the right. **(C-D) Identification of the viral gene segment(s) contributing to host adaptation:** Individual gene segments encoding vRNP from HPhLA D1.1 (LA) were replaced with the viral segments of HPhTX B3.13 (**C**). Individual gene segments encoding vRNP from HPhTX B3.13 (LA) were replaced with the viral segments of HPhLA D1.1 (**D**). **(E) Schematic representation of WT PB2_B3.13 TX (top, red), WT PB2_D1.1 LA (middle, blue) and PB2_D1.1 LA mutants that express either 627K (middle, green) (PB2_D1.1_ 627K) or 701N (bottom, purple) (PB2_D1.1_ 701N). (F) Nluc expression in HEK293T CCSs transfected with the MG, together with the plasmids encoding the polymerase subunits PB2, PB1, PA, and NP of HPhTX B3.13 (TX), HPhLA D1.1 (LA) PB2 WT or PB2 mutants (627K and 701N). (G) Western blot analysis:** Western blot of HEK293T cell lysates transfected as in (**B**), confirming comparable expression of WT PB2 B3.13 TX, PB2 D1.1 LA and mutant proteins (PB2 D1.1_627K and PB2 D1.1_701N). NP served as a transfection control, while empty pCAGGS plasmid-transfected cells were included as negative controls. β-actin was included as a loading control. **(H-M) MG activity at different temperatures:** Nluc and ZsG expressions in HEK293T CCSs transfected with the MG, together with the plasmids encoding the polymerase subunits PB1, PA, NP, and PB2 B3.13 TX or PB2 D1.1 LA, or PB2 mutants at 33^°^C (**H** & **I**, respectively), 37°C (**J** & **K**, respectively) and 39^°^C (**L** & **M**, respectively). **(I, K, M) Fluorescence images:** Representative images of ZsGreen (ZsG) expression were obtained from cells transfected in (**H, J, L**, respectively) using live fluorescence microscopy. Data represent the average of three biological replicates with SD indicated. *p = .01, **p = .001 using unpaired Students t-test. Scale bars = 300 µM.

### Computational interactions between ANP32 and PB2 variants

Following molecular docking (MD) simulations, PBSA calculations were performed on MD-derived snapshots to estimate binding free-energy changes across ANP32A- and ANP32B-containing complexes (**Figs. 6A & 6B**). WT PB2 showed broadly similar median binding energies, whereas the PB2 mutations produced distinct trends for the two host factors. ANP32A complexes retained favorable median binding energies across the mutants, including the 194K+627K double mutant. In contrast, ANP32B complexes showed a progressive shift toward weaker binding with the 627K and 194K+627K PB2 mutants approaching near-neutral median ΔG values. Epistasis analysis indicated near-additive behavior for ANP32A but a non-additive response for ANP32B, where the observed double-mutant effect was smaller than the expected additive penalty (**Fig. 6C**). In addition, the energetic pattern was consistent with the structural behavior observed during MD. RMSD distributions showed relatively limited displacement for most single-mutant complexes, while the 194K+627K ANP32B system displayed the largest spread and highest RMSD values (**Fig. 6D**). Structural overlays further supported this difference, showing that ANP32A remained positioned near the polymerase interface across the mutant systems, whereas ANP32B, particularly in the double-mutant complex, exhibited greater displacement from its initial binding region (**Fig. 6E**). Together, these MD-based structural observations and PBSA estimates suggest that PB2 194K and 627K influence PB2-ANP32 binding in a host-factor-dependent manner, with ANP32A maintaining a more stable association and ANP32B showing reduced binding stability, especially with the double PB2 194K and 627K mutant.

**Figure 6.**
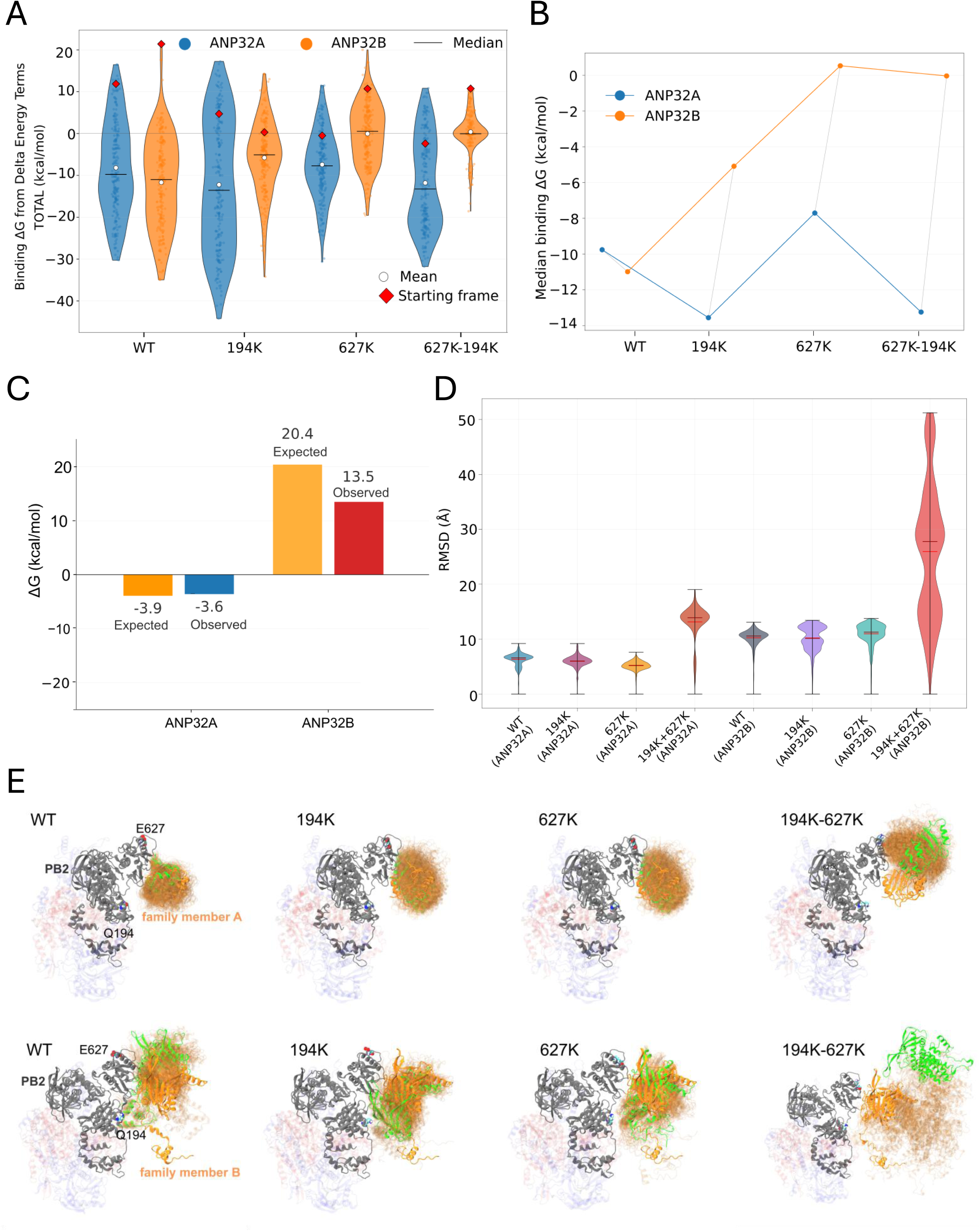
Binding free energy distributions, epistasis, structural dynamics, and host-factor selectivity of PB2 variants with ANP32A and ANP32B. (A-B) Binding free energy distributions were calculated for PB2 WT and mutant complexes with ANP32A and ANP32B using MM/PBSA over 200 trajectory snapshots: The upper-left panel shows binding ΔG distributions for each PB2 variant with ANP32A and ANP32B. Violin plots represent the distribution of binding free energies, small points indicate individual trajectory snapshots, white circles indicate mean values, black horizontal lines indicate medians, and red diamonds indicate starting-frame energies. ANP32A generally maintains favorable binding across variants, with median ΔG values remaining negative, whereas ANP32B shows progressive destabilization, with the double mutant approaching near-zero binding free energy (**A**). The upper-right panel summarizes median binding ΔG values across mutations for ANP32A and ANP32B (**B**). ANP32A retains relatively favorable binding across all variants, with median ΔG values ranging from approximately −7.9 to −13.1 kcal/mol. In contrast, ANP32B exhibits increasing destabilization across the mutant series, with the 194K-627K double mutant approaching near-neutral binding, ΔG ≈ −0.2 kcal/mol, consistent with reduced host-factor compatibility. **(C) Epistatic analysis of the 194K+627K double mutant by comparing expected and observed binding free energy changes:** For ANP32A, the observed value closely matches the expected value, indicating approximately additive behavior: expected ΔG = −3.9 kcal/mol, observed ΔG = −3.6 kcal/mol, deviation = +0.4 kcal/mol, p = 0.858. For ANP32B, the observed destabilization is lower than expected, indicating a significant protective deviation: expected ΔG = +20.4 kcal/mol, observed ΔG = +13.5 kcal/mol, deviation = −7.0 kcal/mol, p < 0.001. **(D) Chain C+D RMSD distributions across all PB2–ANP32 complexes:** ANP32A-associated complexes remain comparatively stable, with RMSD values of WT 6.3 ± 1.1 Å, 194K 6.0 ± 1.0 Å, 627K 5.2 ± 0.8 Å, and 194K+627K 10.2 ± 2.1 Å. ANP32B-associated complexes show higher baseline RMSD values, including WT 13.1 ± 3.1 Å, 194K 10.2 ± 1.5 Å, and 627K 11.0 ± 1.8 Å, with a dramatic increase in the 194K+627K double mutant to 25.9 ± 12.1 Å, indicating severe conformational instability. **(E) Representative structural dynamics for ANP32A and ANP32B complexes across PB2 variants:** PB2 is shown in gray, ANP32 family members are shown with trajectory overlays, and key PB2 residues Q194 and E627 are indicated where visible. Structural overlays progress from early trajectory frames shown in orange/transparent conformations to later/final conformations shown in green. ANP32A variants retain relatively proximal orange and green overlays, consistent with maintained interface remodeling rather than complete dissociation. In contrast, the ANP32B 194K+627K double mutant shows pronounced spatial separation between early orange contact positions and the final green conformation, directly visualizing progressive unbinding and loss of stable host-factor association.

## Discussion

The recent emergence of multiple H5N1 clade 2.3.4.4b genotypes in mammalian hosts has raised important questions regarding the viral determinants that drive mammalian adaptation, pathogenicity, and transmission. Consistent with this concern, several studies have demonstrated that clade 2.3.4.4b HPAIV H5N1 viruses are capable of respiratory droplet transmission in the ferret model, including the 2022 mink-derived isolate from Spain and the 2024 B3.13 bovine- and human-derived isolates from the USA ^12,21,50–52^. In contrast, other studies have reported that both B3.13 and D1.1 human H5N1 isolates failed to transmit via respiratory droplets in ferrets ^53–56^. These conflicting findings indicate that respiratory droplet transmissibility is not a uniform characteristic of clade 2.3.4.4b H5N1 viruses but rather depends on specific viral and potentially host factors. Ferret respiratory droplet transmission efficiency can be influenced by airflow direction, cage design, and other environmental parameters ^57^. Given these inconsistent observations, we evaluated transmission using the direct contact ferret model, a robust approach for assessing mammalian transmissibility while minimizing the variability associated with respiratory droplet transmission experiments of H5N1 viruses. We performed a comprehensive comparison of the first two human isolates of the cattle-associated HPhTX B3.13 genotype and the avian-associated HPhLA D1.1 using *in vitro* systems, ferret infection and transmission studies, viral genetics, polymerase MG assays, and molecular dynamics simulations. Together, our findings demonstrate that the superior pathogenicity and transmissibility of HPhTX B3.13 are primarily associated with enhanced viral polymerase activity rather than differences in HA or NA functions.

One of the most striking observations was the markedly enhanced replication of HPhTX B3.13 in mammalian cells, accompanied by larger plaque morphology and increased viral titers, whereas replication in avian DF-1 cells was comparable between the two genotypes. These findings suggest that the replication advantage of HPhTX B3.13 is host-specific and reflects improved adaptation to mammalian cellular environments rather than an intrinsically faster replicative capacity. Similar host-specific increases in polymerase efficiency have been described during mammalian adaptation of avian influenza viruses, particularly involving mutations within the PB2 segment that facilitate replication in mammalian cells ^38,58^. Despite these pronounced differences in viral genome replication and gene transcription, glycan microarray analyses demonstrated nearly identical receptor-binding profiles for the two viruses, with both retaining a strong preference for avian-type α2,3-linked sialic acid receptors. These observations indicate that enhanced receptor recognition is unlikely to explain the increased replication or transmission of HPhTX B3.13. Instead, they support accumulating evidence that efficient mammalian adaptation of contemporary HPAIV H5N1 can occur without major alterations in HA receptor specificity, provided that post-entry steps of the viral life cycle are optimized. Interestingly, NA analyses revealed that HPhTX B3.13 possessed significantly lower enzymatic activity than HPhLA D1.1 despite exhibiting superior replication. Although this initially appears paradoxical, an optimal functional balance between HA binding and NA cleavage is essential for efficient influenza virus replication ^59,60^. Reduced NA activity may prolong virion attachment at the infected cell surface, thereby promoting localized cell-to-cell spread and contributing to the larger plaque phenotype observed for HPhTX B3.13. Similar observations have been reported for other IAVs, including the 2009 pandemic H1N1 in which partial reductions in NA activity improved viral fitness in humans by maintaining an appropriate HA-NA functional balance ^61,62^. Because both viruses exhibited comparable NA thermostabilities, differences in enzyme stability are unlikely to contribute substantially to the observed phenotypic differences.

The enhanced replication observed *in vitro* translated directly into increased pathogenicity and transmission in ferrets. HPhTX B3.13 infection resulted in greater weight loss, higher clinical scores, earlier mortality, increased viral shedding, broader tissue dissemination, and complete lethality among contact animals. In contrast, HPhLA D1.1 exhibited limited transmission, reduced tissue dissemination, and incomplete infection of exposed contact ferrets. The higher viral loads detected in the respiratory tract and extrapulmonary organs of HPhTX B3.13 likely increased both disease severity and the amount of infectious virus available for transmission. Previous studies have consistently demonstrated that increased viral tropism in the upper respiratory tract is an important determinant of efficient influenza virus transmission ^63,64^, supporting the interpretation that enhanced replication is a fundamental determinant of the increased transmissibility observed for HPhTX B3.13. Histopathological and immunohistochemical analyses further supported these conclusions. Although both viruses infected respiratory and extrapulmonary tissues, HPhTX B3.13 consistently exhibited broader viral antigen distribution and more extensive tissue involvement, particularly within the lungs, brain, trachea, and spleen. The widespread NP antigen detected in contact ferrets infected with HPhTX B3.13 indicates highly efficient transmission leading to systemic dissemination following natural exposure. In contrast, HPhLA D1.1 produced more localized lesions with substantially reduced viral antigen distribution in contact animals, consistent with its lower transmission efficiency. The presence of viral antigen within the central nervous system of HPhTX B3.13 infected ferrets further supports previous observations that highly pathogenic H5N1 viruses possess neuroinvasive potential and suggests that enhanced polymerase activity may facilitate dissemination beyond the respiratory tract ^11,65,66^. The presence of plasma cell-containing infiltrates in the trachea and lungs also motivated future examination of the local B-cell response. A recently established ferret immunoglobulin-gene reference now enables B-cell receptor repertoire sequencing to identify infection-expanded clonotypes which could help to define the antibody responses associated with direct infection and contact transmission ^67^.

Whole-genome sequencing revealed fundamentally different evolutionary trajectories for the two H5N1 genotypes during mammalian infection. HPhTX B3.13 remained genetically stable throughout infection and transmission without evidence of positive selection, suggesting that it is already well adapted for replication in mammals. In contrast, HPhLA D1.1 rapidly acquired the mammalian adaptation PB2 E627K following only a single round of replication in ferrets. Moreover, viruses recovered from contact animals acquired both PB2 E627K and the previously uncharacterized PB2 Q194K substitution. The rapid and repeated emergence of PB2 E627K strongly indicates intense selective pressure for increased polymerase activity during mammalian infection, consistent with decades of influenza evolution studies identifying residue 627 in PB2 as a major determinant of host adaptation ^68–70^. Functional MG assays directly demonstrated that PB2 represents the principal determinant underlying the differences in polymerase activity between these two genotypes. Exchange of PB2 alone largely transferred the high-polymerase phenotype of HPhTX B3.13 to HPhLA D1.1, whereas substitutions of the remaining polymerase components produced substantially smaller effects. These results emphasize the dominant role of PB2 in regulating H5N1 polymerase activity during mammalian adaptation. Importantly, our study identified PB2 Q194K as a previously uncharacterized adaptive mutation that acts synergistically with PB2 E627K to enhance mammalian adaptation. Although PB2 Q194K has been detected previously in conjunction with PB2 T683I during serial guinea pig nasal wash passaging of the mouse-adapted PR8 strain that acquired transmissibility ^71^, its functional contribution has not been completely defined. Our findings provide evidence that PB2 Q194K, together with PB2 E627K, is likely to contribute to increased viral pathogenicity and transmission in ferrets ^12,58^. Whereas PB2 E627K enhanced polymerase activity at all the temperatures examined, PB2 Q194K alone increased polymerase activity only at 37°C and 39°C but not at 33°C. Because the upper respiratory tract of ferrets is maintained near 33°C, the inability of PB2 Q194K alone to enhance polymerase activity at this temperature may explain why HPhLA D1.1 retains limited viral shedding and inefficient transmission despite acquiring this mutation. In contrast, the combined PB2 E627K/Q194K mutant restored polymerase activity to levels approaching those of HPhTX B3.13 across all tested temperatures, suggesting that these mutations cooperate to optimize polymerase function throughout the mammalian respiratory tract. These findings identify PB2 Q194K as a potential secondary adaptive mutation that enhances the effects of PB2 E627K during mammalian adaptation.

Our computational analyses provide a possible structural explanation for these functional observations. Molecular dynamics simulations demonstrated that PB2 mutations differentially affect interactions with host ANP32 proteins, which are essential co-factors for influenza polymerase activity ^38–40^. While ANP32A complexes remained relatively stable following the introduction of PB2 E627K and PB2 Q194K, ANP32B complexes exhibited progressively weaker binding and increased structural flexibility, particularly in the double-mutant complex. These results suggest that mammalian-adaptive PB2 mutations may alter polymerase–host interactions in a host-factor-dependent manner rather than simply by strengthening binding affinity. Such dynamic modulation of ANP32 utilization may contribute to enhanced polymerase efficiency during mammalian infection while preserving sufficient structural flexibility for optimal viral replication. The finding that currently available CDC CVVs generate broadly cross-neutralizing antibodies against both HPhTX B3.13 and HPhLA D1.1 provides encouraging evidence that these emerging genotypes remain antigenically similar despite their marked biological differences. Thus, increased virulence and transmission appear to result primarily from enhanced replication fitness rather than substantial antigenic drift.

Overall, our study supports a model in which mammalian adaptation of contemporary HPAIV H5N1 is driven predominantly by optimization of viral polymerase function rather than by changes in receptor specificity. The genetic stability of HPhTX B3.13, together with its high polymerase activity, efficient transmission, and increased pathogenicity, suggest that this genotype is already well adapted for mammalian replication. In contrast, HPhLA D1.1 remains incompletely adapted but rapidly acquires mammalian-adaptive PB2 mutations during replication in ferrets. The identification of PB2 Q194K as a cooperative mutation that enhances the activity of PB2 E627K expands our understanding of influenza polymerase adaptation and identifies a potential molecular marker for the surveillance of emerging H5N1 viruses with increased zoonotic potential.

Finally, this study has some limitations. First, transmission was assessed in a single mammalian model with a relatively small sample size under controlled laboratory conditions, which may not fully recapitulate natural exposure settings or host diversity. Second, only one isolate per genotype was evaluated. Third, the potential contribution of sex to pathogenicity or transmission was not assessed since only female ferrets were used in this study and because some previous studies used male ferrets ^17,72^. Finally, our *in silico* computational analysis suggested that the PB2 mutations 194K and 627K affect PB2-ANP32 binding in a host-dependent manner, with APN32A maintaining a more stable association than ANP32B and showing reduced binding stability. However, future studies are needed to confirm this hypothesis.

## Declarations

## Acknowledgments

We thank the Histology Unit Team staff, Dr. Renee Escalona and Mr. Colin Chuba, for their assistance in tissue staining/immunostaining experiments, and the Cell Biology Core Lab at Texas Biomed for assistance with the multiplex cytokine assay. We thank Ashley Gay-Cobb for the technical support. We acknowledge BEI Resources for providing PB1 (Clone F5-46) and PA (Clone 1F6) MAbs and N1 influenza A/dairy cattle/Texas/24-008749-001-original/2024 H5N1 recombinant protein (NR-59817). We thank Dr. Daniel Perez at the Animal Health Research Center, Center for Vaccines and Immunology, Department of Population Health, Poultry Diagnostic and Research Center for providing MDBK cells. We also thank Drs. Han Di and Bin Zhou at the Center for Disease Control and Prevention (CDC) for providing the ferret antisera samples IDCDC-RG71A, IDCDC-RG78A, and IDCDC-RG80A.

## Authors Contributions

Conceptualization: A.M.E., G.C.I., and L.M-S.; Methodology: A.M.E., R.S.B., M.B., A.P., H.B., V.S., R.N.P., F.B., J.C., A.R., J.L., C.Y., A.N., R.P.V. and E.M.A.; Data collection and interpretation: A.M.E, M.B., R.N., T.J.C.A., R.P.V., G-J.B., A.G-S., E.M.A., G.C.I. and L.M-S.; Funding acquisition and resources: A.M.E., A.G-S., G.C.I., E.M.A. and L.M-S.; Writing-original draft preparation: A.M.E. and L.M-S.; Writing—review and editing: all authors have read and agreed to the published version of the manuscript.

---

**Supplementary Fig. S1.**
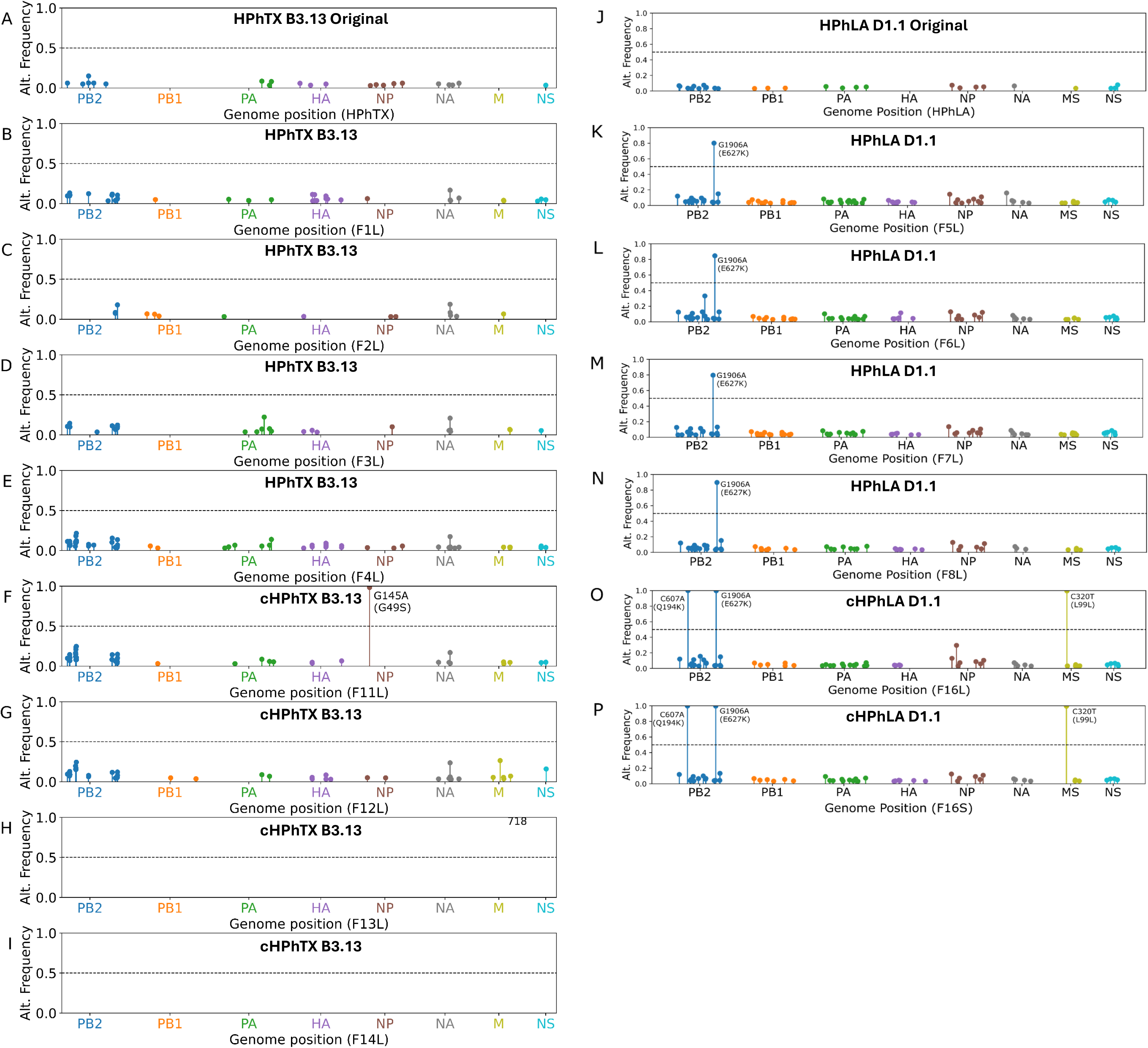
NGS analysis of viral adaptation in ferrets: Viral genomes were sequenced from the original HPhTX B3.13 inoculum (**A**), tissue homogenates from HPhTX B3.13 infected ferrets (**B–E**), and HPhTX B3.13 contact ferrets (**F–I**). Viral genomes were also sequenced from the original HPhLA D1.1 inoculum (**J**), tissue homogenates from HPhLA D1.1 infected ferrets (**K–N**), and HPhLA D1.1 contact ferrets (**O–P**). **FxL** or **FxS** on the x-axis denotes the ferret identification number (**x**) and the tissue analyzed, where **L** and **S** represent lung and spleen tissue homogenates, respectively, from which viral RNA was extracted for next-generation sequencing (NGS). Nucleotide substitutions are indicated together with their corresponding amino acid changes (in parentheses).

**Supplementary Fig. 2.**
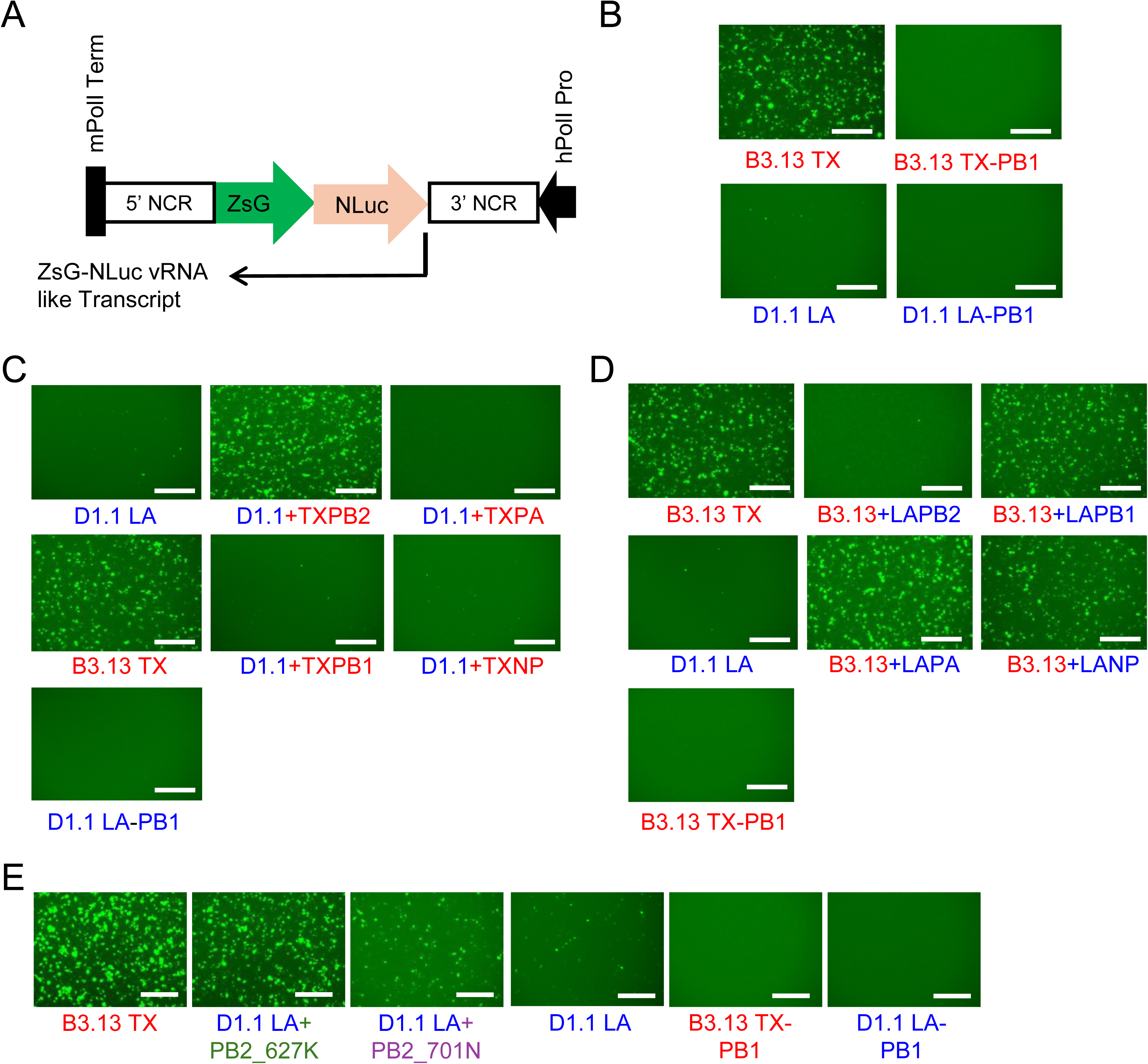
ZsGreen (ZsG) expression showing enhanced polymerase activity of HPhTX B3.13 (TX) and HPhLA D1.1 mutants 627K and 701N compared to HPhLA D1.1 WT. (A) Schematic representation of the H5N1 MG plasmid: The human polymerase I promoter (hPolI Pro) is indicated as a black arrow. The mouse polymerase I terminator (mPolI Term) is indicated as a black box on the opposite side. 5’ and 3’ Non-coding regions (NCR) of the H5N1 NP segment are indicated with white boxes. The ZsG fused to Nanoluciferase (Nluc) to express a viral-like transcript is also indicated. **(B) Representative images of ZsG expression from cells transfected in Fig 5A using live fluorescence microscopy. (C) Representative images of ZsG expression from cells transfected in Fig 5C using live fluorescence microscopy. (D) Representative images of ZsG expression from cells transfected in Fig 5D using live fluorescence microscopy. (E) Representative images of ZsG expression from cells transfected in Fig 5E using live fluorescence microscopy.** Scale bars = 300 µM.

**Supp table 1.** Amino Acid variations between HPhTX B3.13 and HPhLA D1.1.

| Protein | Residue | HPhTX B3.13 | HPhLA D1.1 |
| --- | --- | --- | --- |
| PB2 | 58 | A | T |
|  | 109 | I | V |
|  | 139 | I | V |
|  | 441 | N | D |
|  | 495 | I | V |
|  | 627 | K* | E |
|  | 649 | I | V |
|  | 676 | A | T |
| PB1 | 16 | N | D |
|  | 59 | S | T |
|  | 75 | D | E |
|  | 154 | G | S |
|  | 171 | V | M |
|  | 172 | E | D |
|  | 179 | I | M |
|  | 207 | K | R |
|  | 375 | S | N |
|  | 587 | P | A |
|  | 614 | E | D |
|  | 694 | N | S |
| PB1-F2 | 4 | E | G |
|  | 7 | I | T |
|  | 8 | P | Q |
|  | 12 | L | S |
|  | 18 | I | T |
|  | 20 | K | R |
|  | 21 | K | R |
|  | 22 | G | E |
|  | 31 | G | E |
|  | 36 | I | T |
|  | 40 | D | G |
|  | 42 | C | Y |
|  | 44 | M | R |
|  | 46 | M | T |
|  | 47 | S | N |
|  | 49 | V | A |
|  | 54 | R | Q |
|  | 55 | T | I |
|  | 57 | S | C |
|  | 58 | L | W |
|  | 65 | K | R |
|  | 66 | N | S |
|  | 68 | I | T |
|  | 70 | E | G |
|  | 75 | R | L |
|  | 82 | L | S |
|  | 84 | N | S |
|  | 90 | S | N |
| PA | 61 | M | I |
|  | 85 | A | T |
|  | 113 | R | K |
|  | 269 | R | K |
|  | 277 | P | S |
|  | 322 | I | V |
|  | 323 | V | I |
|  | 348 | I | L |
|  | 388 | S | G |
|  | 391 | R | K |
|  | 400 | S | P |
|  | 441 | V | M |
|  | 545 | I | V |
|  | 558 | L | S |
|  | 608 | S | T |
|  | 626 | K | R |
| **PA-X** | 61 | M | I |
|  | 85 | A | T |
|  | 113 | R | K |
|  | 193 | N | S |
|  | 195 | R | K |
|  | 219 | F | S |
|  | 250 | Q | P |
|  | 252 | R | K |
| **HA** | 11 | V | I |
|  | 52 | T | A |
|  | 120 | M | L |
|  | 131 | Q | L |
|  | 211 | I | T |
|  | 226 | A | V** |
|  | 341 | K | R |
|  | 491 | N | D |
|  | 526 | V | I |
| **NP** | 52 | H | Y |
|  | 482 | N | S |
| **NA** | 8 | T | I |
|  | 20 | V | I |
|  | 23 | M | V |
|  | 44 | Y | N |
|  | 45 | Q | H |
|  | 48 | P | T |
|  | 53 | I | V |
|  | 74 | F | L |
|  | 75 | L | I |
|  | 81 | T | D |
|  | 82 | S | P |
|  | 84 | T | A |
|  | 221 | N | S |
|  | 234 | V | I |
|  | 241 | V | I |
|  | 257 | K | R |
|  | 269 | M | L |
|  | 286 | G | S |
|  | 287 | D | E |
|  | 288 | I | V |
|  | 321 | I | V |
|  | 329 | N | S |
|  | 336 | S | G |
|  | 338 | M | V |
|  | 339 | P | S |
|  | 395 | E | A |
| **M1** | 85 | S | N |
|  | 87 | T | N |
|  | 200 | V | A |
|  | 227 | T | A |
| **M2** | 61 | G | R |
|  | 88 | N | D |
| **NS1** | 7 | L | S |
|  | 75 | E | G |
|  | 83 | S | P |
|  | 87 | P | S |
|  | 116 | S | C |
|  | 139 | N | D |
|  | 147 | L | I |
|  | 171 | D | N |
|  | 193 | R | Q |
|  | 223 | E | A |
| **NEP** | 7 | L | S |
|  | 36 | E | K |
|  | 63 | G | E |

\* E627K allows avian-origin viruses to evade host restrictions, drastically boosting polymerase activity, viral replication, and pathogenicity in mammals (1, 2).
\*\* A226V affects receptor binding affinity and binding to horse RBCs (3).

